# Copy number variation of the putative speciation genes in the European house mouse hybrid zone

**DOI:** 10.1101/2021.12.22.473661

**Authors:** Alexey Yanchukov, Zuzana Hiadlovská, Zeljka Pezer, Miloš Macholán, Jaroslav Piálek, Stuart J.E. Baird

## Abstract

Hybrid zones have long been described as “windows on the evolutionary process”, and studying them has become even more important since the advance in the genome analysis tools. The hybrid zone between two subspecies of the house mouse (*Mus musculus musculus* and *Mus m. domesticus*) is a unique model speciation system to study fine scale interactions of recently diverged genomes. Here, we explore the role of gene Copy Number Variation in shaping the barrier to introgression in the hybrid zone within a previously established transect in Central Europe. The CNV of seven pre-selected candidate genes was determined via droplet-digital PCR and analyzed in the context of ~500k SNPs, with the ancestral population (i.e. *musculus* or *domesticus*) of every SNP allele previously inferred in the admixed individuals (Baird et al., *in prep*.). The copy numbers of five genes were clearly associated with the prevalence of either *musculus* or *domesticus* genomes across the hybrid zone. In three cases, the highest and/or outlying levels of association were observed at or very close to the annotated positions of the respective gene amplicons, demonstrating the power of our approach in confirming the reference locations of copy number variants. Notably, several other reference locations were recognized as positive outliers in the association with particular CNV genes, possibly representing the extra gene copies and/or their epistatic interaction sites.

## Introduction

The genome-wide copy number variation (CNV) observed among eukaryote genomes is an extensive, ubiquitous and complex phenomenon (Redon et al. 2006; Pirooznia, Goes, and Zandi 2015). Of particular interest are the findings that CNV contributes to differentiation between recently diverged populations and that genes mapped to CNV regions are involved in adaptation to new environments (Bryk et al. 2013; Bryk and Tautz 2014; Perry et al. 2007; Chain et al. 2014; Mejía-Benítez et al. 2015; Iskow, Gokcumen, and Lee 2012), species-specific mate recognition (Emes et al. 2004; C. M. Smadja et al. 2015; Paudel et al. 2015), and immune response (Ottolini et al. 2014; Roberts et al. 2014), i.e. the processes known to be important in the course of speciation. Among a large number of such studies, however, only a few were designed to investigate the role of CNV in speciation as a primary goal (Locke et al. 2015; Chain et al. 2014). One reason for that is a need for genome-wide CNV scans across a considerable number of specimens from naturally evolving populations, conditionally restricted by the fact that the quality of such scans decreases as the target specimen’s genome diverges from the annotated genome assembly of that particular organism (i.e. suffers from ascertainment bias, Boursot and Belkhir 2006). This complicates genotyping individuals from distant populations or from other species. Also, genome-wide studies typically have modest sample sizes so drawing conclusions about the population dynamics of particular CNVRs is often difficult. At the same time, much is known about the speciation genes, at least in the model vertebrate systems, to warrant a candidate gene approach in the CNV speciation studies. Genotyping of a few CNV genes can be easily performed on large samples and allows flexibility in experimental design to correct for ascertainment bias, thus prompting clearer results. Indeed, the famous evidence for the role of CNV in adaptation and population differentiation comes from a candidate gene study aimed at the salivary amylase gene *AMY2* where a higher copy number has likely evolved in response to specialization for starch-rich diet in several human populations (Perry et al. 2007; Mejía-Benítez et al. 2015). Deep insight into the role of CNV in adaptation is possible through combining the CNV data with the results on expression, function and evolutionary history of the candidate genes (Iskow, Gokcumen, and Lee 2012; Henrichsen et al. 2009; Conrad et al. 2010).

The question of why CNV are predisposed for adaptation and speciation is mostly addressed by the theoretical works. It is predicted that genes existing in multiple copies could be readily recruited for adaptation via (i) direct selection for increased gene dosage (Kondrashov and Koonin 2004; F. a. Kondrashov 2012; Cardoso-Moreira et al. 2016) and (ii) functional redundancy which allows the multiple copies to specialize without compromising the existing gene function (Michael Lynch and Force 2000; M. Lynch 2000). The opportunities for adaptive response increase in dynamic ecological (Kondrashov 2012; Katju and Bergthorsson 2013) and genetic (Yeaman and Whitlock 2011; Yeaman and Otto 2011; Otto and Yong 2002) contexts. In nature, both of these contexts are abound in zones of secondary contact between recently diverged populations (hybrid zones), which harbor large numbers of highly recombinant genomes and can therefore create favorable environment for evolution of multicopy gene systems (Yanchukov and Proulx 2012; Yanchukov and Proulx 2014). In this paper, we make the first step towards resolving CNV in genes putatively linked to the speciation process, by looking at the candidate gene copy number genotypes combined with an existing dense SNP dataset covering the hybrid zone and the allopatric populations of the European house mouse (*Mus musculus*).

The Eastern European house mouse, *M. m. musculus*, and the Western European house mouse, *M. m. domesticus*, meet and hybridize along > 2500 km long narrow contact zone running from Norway to the Black Sea (Jones et al. 2010; Baird & Macholán 2012; Ďureje et al. 2012). The house mouse hybrid zone (HMHZ) has been studied extensively across multiple transects and benefited from the powerful arsenal of molecular and genomic tools developed for the house mouse as primary model for biomedical and evolutionary research (Macholán et al. 2012; Phifer-Rixey and Nachman 2015). Introgression patterns of allozyme, microsatellite and SNP markers along with the experimental work on hybrid fitness (Albrechtová et al. 2012; Turner et al. 2012, 2014) suggest that the zone is maintained by the balance between selection against hybrids / inter-subspecific crosses and gene flow, i.e. is an example of a tension zone (Gay et al. 2008;Barton and Hewitt 1985). Multiple genomic regions (overrepresented on the X chromosome) are responsible for genetic incompatibilities between the two subspecies (Payseur et al. 2004; Harr 2006; Macholán, Munclinger, Sugerková, et al. 2007; Dufková et al. 2011, Turner, Schwahn, and Harr 2012; Janousek et al. 2012). Preferential gamete recognition within subspecies might contribute to pre-zygotic isolation between *musculus* and *domesticus* (Dean and Nachman 2009; but see Firman et al. 2014) and decreased sperm number/function was found to be a major factor contributing to selection against hybrids ( Good et al. 2008; Albrechtová et al. 2012; Turner, Schwahn, and Harr 2012). Behavioral isolation, as evident from the clinal variation of the urinary and salivary signal protein genes and from behavioral preference tests for these signals, may also play a role in maintaining the barrier between the two subspecies (Vošlajerová Bímová et al. 2009, 2011; Talley, Laukaitis, and Karn 2007; C. M. Smadja et al. 2015).

The wide spectrum of isolation mechanisms is reflected by similarly complex picture of genomic admixture in HMHZ. The most recent analysis of a set of high-resolution SNP markers genotyped on the Affymetrix Mouse Diversity Array (MDA, Yang et al. 2009) across a part of HMHZ on the border between western Bohemia (Czech Republic) and north-eastern Bavaria (Germany) allowed for mapping the contributions of the *M*. *m. musculus* and *M*. *m. domesticus* ancestral populations in the genomes of hybrids at high resolution (approximately one diagnostic SNP per 500 kb, Baird et al., *in prep*.). Large-scale introgression, i.e. the presence in predominantly *musculus* or *domesticus* genomes of multiple wide regions that trace their ancestry to *domesticus* or *musculus* populations, respectively, was found on every chromosome except the X (Baird et al. *in prep*.). This pattern could have resulted from selection against gene flow at some regions and/or favoring it at the others: blocks of neighboring SNPs would then show a similar pattern depending on the strength of physical (map distance) or epistatic association with selected loci (Barton and Bengtsson 1986; Barton 1983; Baird 1995). The high-resolution SNP maps can thus be used to *a priori* assess the relative importance of particular genomic regions in divergence and speciation in *musculus* – *domesticus* system. Comprehensive gene and sequence ontology annotation of the introgressed regions can provide additional clues pointing to the loci potentially relevant in speciation (Baird et al., *in prep*.).

Two recent genome-wide scans for structural variation among a few geographically distant populations of *M. m. domesticus* reveal that (i) CNV pattern of ~1900 transcribed sequences reflects the history of neutral population divergence and (ii) previously reported regions of segmental duplications are significantly overrepresented as CNV (Bryk and Tautz 2014; Pezer et al. 2015). Independent analysis of the hybridization probe signal intensity data of 351 (mostly classical laboratory strains) mouse specimens confirms the latter finding (Locke et al. 2015). These results suggest that recurrent CNV regions (CNVR) are very common in the house mouse, and that the same CNVRs are likely to show up in previously unstudied populations. Furthermore, greater standing variation should preferentially expose such loci to natural selection, thus setting up the most likely trajectory for the adaptation process. Indeed, the three environment-responsive genes that also have the largest differences in their copy numbers among three *M. m. domesticus* populations (Pezer et al. 2015) have all been previously reported as CNVs by at least four independent studies (Keane et al. 2011; Henrichsen et al. 2009; Wong et al. 2012; Quinlan et al. 2010). In short, genotyping of recurrent CNVRs should narrow down the genomic target in search of regions involved in speciation.

The previous advances in hybrid zone and CNV research combined with a rich spectrum of molecular resources available for the house mouse make HMHZ an ideal system for a pilot empirical exploration of the role of CNV in house mouse speciation process. In this paper, we genotype the copy numbers of several of candidate genes that are likely to be involved in both CNV and speciation. We then directly use the results of a parallel analysis of the MDA data (Baird et al., *in prep*.) to explore the genome-wide association of the CNV types.

## Results

We compiled a list of all protein coding genes from the Mouse Genome Database (Eppig et al. 2015, http://www.informatics.jax.org) that are also completely contained within the known mouse CNVRs, accessible in the NCBI’s database of genomic structural variation (dbVar, Lappalainen et al. 2013) in June-July 2014. This search yielded ~# genes. We then narrowed our search to those genes that (i) are reported to be copy number variable by at least three independent studies, (ii) possess a known biological function potentially related to speciation (iii) located within or close to the introgressed regions of the house mouse genome in HMHZ (Baird et al. *in prep*) and (iv) allow for design of specific primers each with at least 3 basepair mismatches to the next most likely target amplicon. Many members of large gene families did not satisfy the amplification specificity criterion (iv), further narrowing the list down to a few tens of candidate genes (data not shown).

According to the house mouse reference genome assembly CRCm38 (mm10) in NCBI’s dbVar (Lappalainen et al. 2013), four CNV amplicons in our study are located uniquely within the coding regions of the following genes: mast cell chymase 1 (*Cma1*), sperm motility kinase 2a (*Smok2a*), spermatogenesis associated glutamate (E)-rich protein 4A (*Speer4a*) and defensin beta 7 (*Defb7*). One amplicon has a 100% shared similarity to two interferon-induced proteins with tetratricopeptide repeats 3 (*Ifit3*) and 3b (*Ifit3b*). These genes are separated by 19.2 kb and are consistently reported in the same continuous CNVR by 3 independent studies (Henrichsen et al. 2009; Keane et al. 2011; Quinlan et al. 2010). Finally, two amplicons are located in the short intergenic regions between vomeronasal type-2 receptors 5 and 6 (*Vmn2r5* and *Vmn2r6*), and between receptors 6 and 7 (*Vmn2r6* and *Vmn2r7*). These amplicons had been mapped to either one (Cutler et al. 2007; Pezer et al. 2015) or two (Keane et al. 2011) distinct CNVRs. In the following, we retain the gene names for unique amplicons (e.g. *Cma1* for *Cma1* gene), but use the abbreviations *Ifit3|3b* to indicate the amplicon for *Ifit3* and *Ifit3b*, and *Vmn2r5:6, Vmn2r6:7* for the amplicons between the corresponding vomeronasal receptor genes. The detailed information about the ddPCR assays and amplification targets used in this study is summarized in Suppl Table 1. For the purpose of our study, we use the terms “CNV gene” and “CNV amplicon” interchangeably.

### CNV in the allopatric musculus and domesticus samples

We designate all geographically distant samples east and north of the hybrid zone as the allopatric *M. m. musculus* samples, and those west and south of the hybrid zone as the allopatric *M. m. domesticus* samples (with the exception of distant mixed populations marked in green on Figure 1A). This division necessarily simplifies the complex evolutionary histories of multiple geographically distant populations, nevertheless the heterogeneity within each of these broad geographic areas has been shown to be much lower than the differences between the respective subspecies (Rajabi-Maham, Orth, and Bonhomme 2008; Hardouin et al. 2015). The hybrid index (*HI*) in the *domesticus* allopatric samples ranged between 0.922 and 0.999, and in the *musculus* allopatric samples between 0.7*10^-3^ and 0.06 (Baird et al., *in prep*.). We found significant differences in mean copy numbers in *Cma1, Smok2a* and *Speer4a* genes, with higher mean CN in *musculus* in each case. The mean CNs of the remaining genes were similar in both subspecies (Table 1). The *V_ST_* statistic, i.e. the relative measure of between-population versus within-population differentiation in gene copy number (Redon et al. 2006, see Material and Methods), ranged from the maximum of 0.83 for *Cma1* to −0.03 for *Vmn2r5:6* (Table 1).

**Figure 1.**
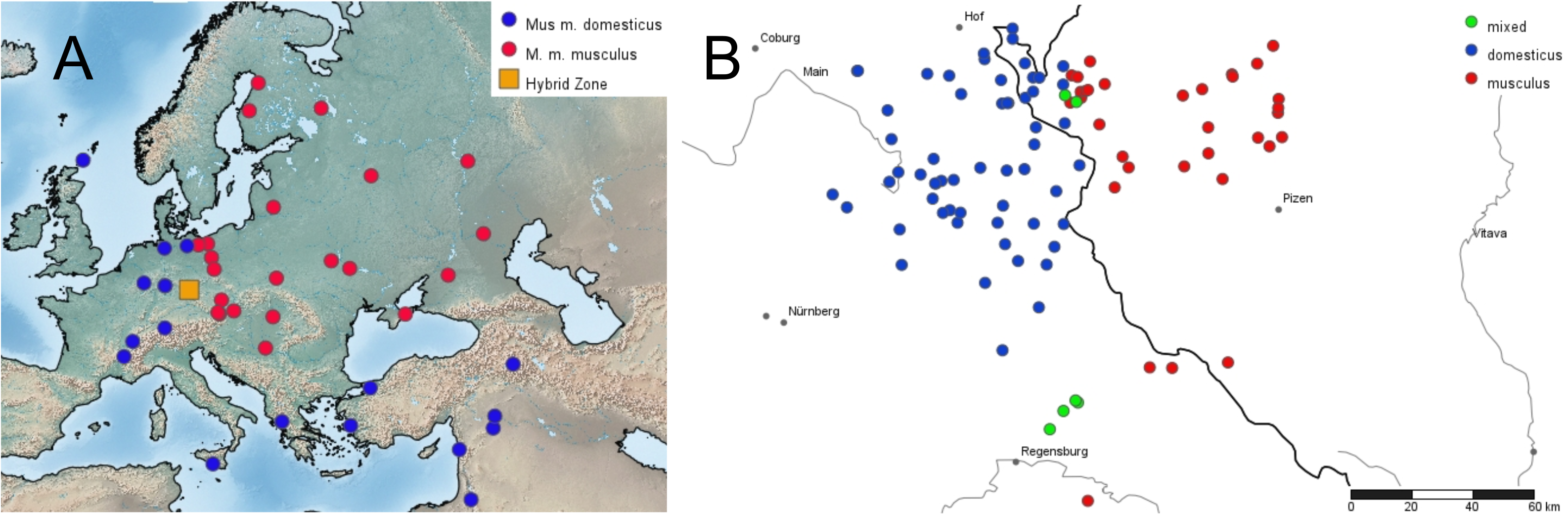
House mouse sampling locations used for CNV genotyping. **A** – Allopatric samples across Europe and Near East. *M. m. domesticus* and *M. m. musculus* populations are shown by blue and red dots, respectively, and the location of the Czech-Bavarian HMHZ transect is indicated by the orange square. One sample from Kazakhstan is outside the map range and is not shown. **B** – zoom in view of the sampling localities across HMHZ: predominantly *domesticus* samples with the average *HI* > 0.7 are shown by blue and predominantly *musculus*samples with *HI* > 0.3 are shown by the red dots, respectively. A few samples with 0.3 < *HI* < 0.7 are in green.

**Table 1.**
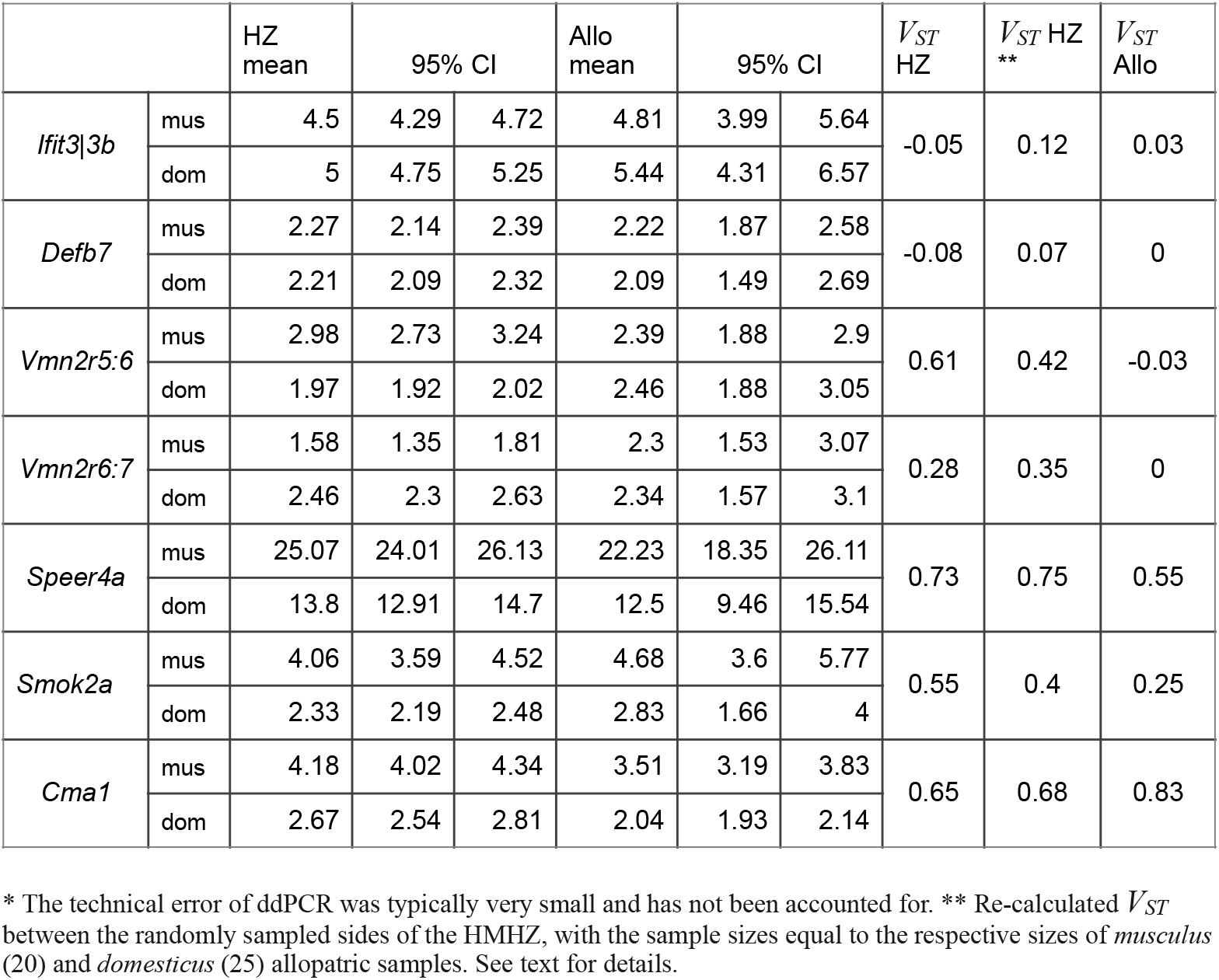
Summary and *V_ST_* statistics* in the allopatric samples and the sides of HMHZ

| | | HZ<br>mean | 95% CI | | Allo<br>mean | 95% CI | | $V_{ST}$<br>HZ | $V_{ST}$ HZ<br>** | $V_{ST}$<br>Allo |
| --- | --- | --- | --- | --- | --- | --- | --- | --- | --- | --- |
| *Ifit3\|3b* | mus | 4.5 | 4.29 | 4.72 | 4.81 | 3.99 | 5.64 | -0.05 | 0.12 | 0.03 |
|  | dom | 5 | 4.75 | 5.25 | 5.44 | 4.31 | 6.57 |  |  |  |
| *Defb7* | mus | 2.27 | 2.14 | 2.39 | 2.22 | 1.87 | 2.58 | -0.08 | 0.07 | 0 |
|  | dom | 2.21 | 2.09 | 2.32 | 2.09 | 1.49 | 2.69 |  |  |  |
| *Vmn2r5:6* | mus | 2.98 | 2.73 | 3.24 | 2.39 | 1.88 | 2.9 | 0.61 | 0.42 | -0.03 |
|  | dom | 1.97 | 1.92 | 2.02 | 2.46 | 1.88 | 3.05 |  |  |  |
| *Vmn2r6:7* | mus | 1.58 | 1.35 | 1.81 | 2.3 | 1.53 | 3.07 | 0.28 | 0.35 | 0 |
|  | dom | 2.46 | 2.3 | 2.63 | 2.34 | 1.57 | 3.1 |  |  |  |
| *Speer4a* | mus | 25.07 | 24.01 | 26.13 | 22.23 | 18.35 | 26.11 | 0.73 | 0.75 | 0.55 |
|  | dom | 13.8 | 12.91 | 14.7 | 12.5 | 9.46 | 15.54 |  |  |  |
| *Smok2a* | mus | 4.06 | 3.59 | 4.52 | 4.68 | 3.6 | 5.77 | 0.55 | 0.4 | 0.25 |
|  | dom | 2.33 | 2.19 | 2.48 | 2.83 | 1.66 | 4 |  |  |  |
| *Cma1* | mus | 4.18 | 4.02 | 4.34 | 3.51 | 3.19 | 3.83 | 0.65 | 0.68 | 0.83 |
|  | dom | 2.67 | 2.54 | 2.81 | 2.04 | 1.93 | 2.14 |  |  |  |

\* The technical error of ddPCR was typically very small and has not been accounted for. \*\* Re-calculated $V_{ST}$ between the randomly sampled sides of the HMMZ, with the sample sizes equal to the respective sizes of *musculus* (20) and *domesticus* (25) allopatric samples. See text for details.

### CNV on the musculus and domesticus sides of the hybrid zone

The distribution of *HIs* among the 207 HMHZ mice is bimodal, with most individuals carrying large relative proportions of either *musculus* or *domesticus* SNPs and only a few with HI in between 0.3 and 0.7 (Baird at al. *in prep*.). It is therefore reasonable to divide the dataset into the *musculus* (i.e. with *HI* < 0.5) and *domesticus* (*HI* > 0.5) sides of the hybrid zone, which also reflects geographic distribution of the sampling localities (Figure 1). The distribution of the mean CNs and the per-gene *V_ST_* statistic between the sides of the HMHZ corresponds generally to that between the respective allopatric samples (Table 1, Figure 2). An exception are *Vmn2r5:6* and *Vmn2r6:7* genes, which both show almost no differentiation between the allopatric samples (*V_ST_* ~ 0), but strong differentiation between the sides of HMHZ (CI bins for *V_ST_* = [0.54, 0.66] and [0.22, 0.32], respectively, see Table 1 for means). To demonstrate that this result cannot be caused by the different sample sizes in the two *V_ST_* comparisons (i.e. more individuals sampled on the sides of HMHZ than in allopatry), we chose randomly the equivalent numbers of mice from the sides of HMHZ as in two allopatric samples and re-calculated *V_ST_* repeatedly over ~1000 iterations. The average *V_ST_* values at *Vmn2r5:6* and *Vmn2r6:7* remained high (*V_ST_* = 0.42 and 0.35, respectively, Table 1).

**Figure 2.**
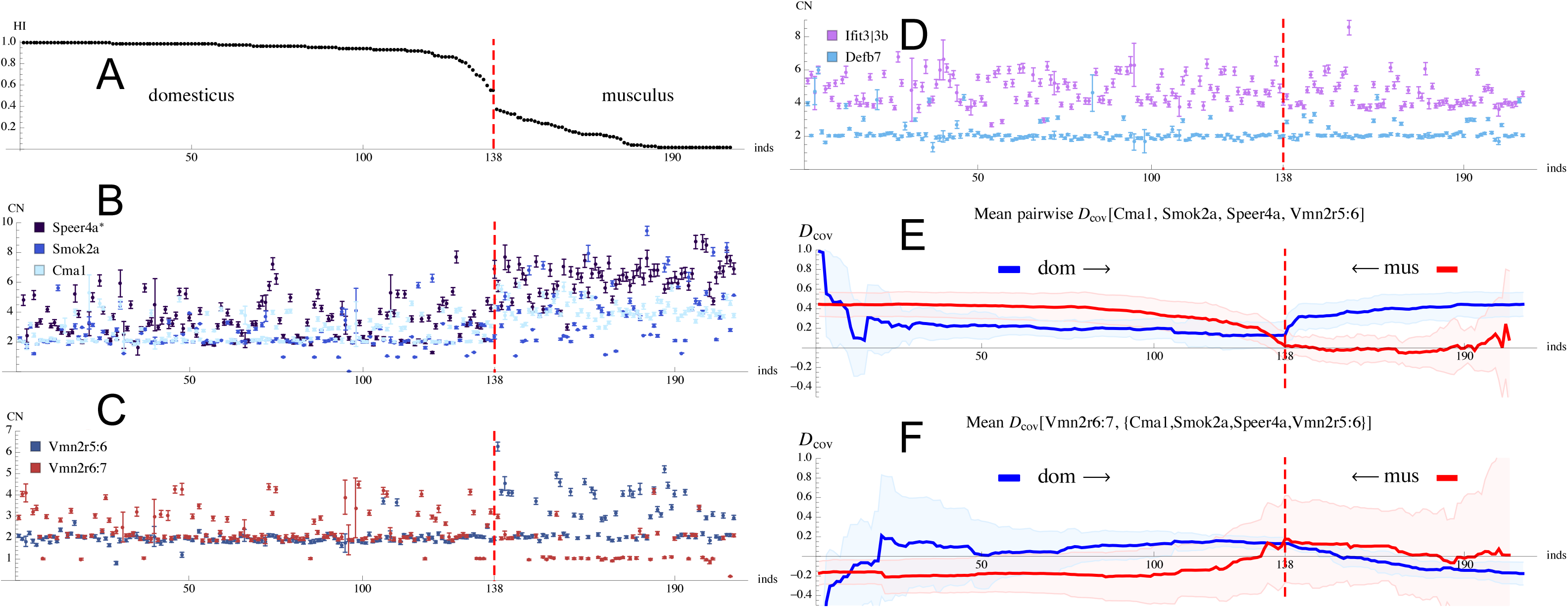
Genome differentiation and copy number variation across HMHZ. **A** – Individual Hybrid Index (*HI*) in descending order from *domesticus* (*HI* = 1) to *musculus* (*H_t_* = 0). The red dotted line is plotted at *HI* = 0.5. **B** – Individual Copy Numbers (CN) of *Speer4a*, *Smok2a* and *Cma1* genes, ordered by the hybrid index. The vertical error bars indicate the standard measurement error (SE) provided by the ddPCR instrument. C – same for *Vmn2r5:6* and *Vmn2r6:7* genes, **D** – same for *Ifit3|3d* and *Defb7* genes. **E** – mean pairwise covariance of gene copy numbers, *D*_cov_ calculated cumulatively, i.e. by adding the individuals one-by-one, from *domesticus* to *musculus* (blue line) and from *musculus* to *domesticus* (red line) on the set of HMHZ individuals ordered by the hybrid index. The following genes were included: *Cma1*, *Smok2a*, *Speer4a* and *Vmn2r5:6*. Shaded colored areas indicate 95% confidence intervals for the mean *D*_cov_. **F** – the same as in **E**, but *D*_cov_ is averaged over pairs where the first member of the pair is *Cma1, Smok2a, Speer4a* and *Vmn2r5:6*, respectively, and the second is *Vmn2r6:7*.

At this stage, notable differences became apparent among the seven CNV genes genotyped in our study. The copy number variation in the group of five genes clearly distinguished the house mouse subspecies in both allopatry and HMHZ (*Cma1*, *Speer4a*, *Smok2a*), or only in HMHZ (*Vmn2r5:6* and *Vmn2r6:7*). We will therefore refer to these five CNV genes as “diagnostic”, as opposed to the “non-diagnostic” genes *Ifit3|3b* and *Defb7* that display no obvious pattern of differentiation between *musculus* and *domesticus* in the allopatry nor in HMHZ.

#### Covariance of gene copy numbers

Strong associations between seemingly unrelated traits and genetic markers are observed in most hybrid zones (N. H. Barton and Hewitt 2003), including HMHZ (Macholán et al. 2008; Macholán et al. 2011; Baird et al., *in prep*.). The primary cause for this phenomenon is selection acting indiscriminately against large segments of genome, before they are able to recombine away from the loci responsible for decreased fitness of hybrids (N. H. Barton 1983). The resulting linkage disequilibria between the underlying alleles derived from one or the other hybridizing taxon cause an increase in covariance among the quantitative traits in the center of the hybrid zone, provided that the mean trait values are different between the taxa (Nürnberger et al. 1995). On the other hand, strong barrier to gene flow will likely maintain similarly strong associations across the entire hybrid zone regardless of proximity to its center (Kruuk et al. 1999).

Since we infer the copy numbers from the ddPCR “phenotypes”, formed by the joint action of multiple gene copies, our CNV data can be treated similarly to polygenic quantitative traits. We therefore calculated the pairwise normalized covariance, *D*_cov_, between the copy number genotypes of the diagnostic genes (Figure 2,3). The covariance was estimated separately on the HMHZ sides and within the allopatric samples (Figure 3). First, the overall covariance was much stronger in the *musculus* allopatric sample compared to either *domesticus* allopatric samples or to both sides of the HMHZ (Fig 3). A distinct pattern was observed for *Vmn2r5:6* gene, which displayed weak negative covariance in all gene pairs in *domesticus* allopatric samples, while the covariance in *musculus* allopatric samples was uniformly positive (Figure 3). The most notable pattern was revealed for *Vmn2r6:7* when both sides of HMHZ were pooled together. Unlike the rest of the genes, the *Vmn2r6:7* copy numbers displayed negative covariance in all pairs (min *D*_cov_ = −0.23). Recall that *Vmn2r6:7* is represented by a higher number of copies on the *domesticus* side of the HMHZ, unlike the rest of the diagnostic genes that on average have more copies on the *musculus* side of the HMHZ (Table 1).

**Figure 3.**
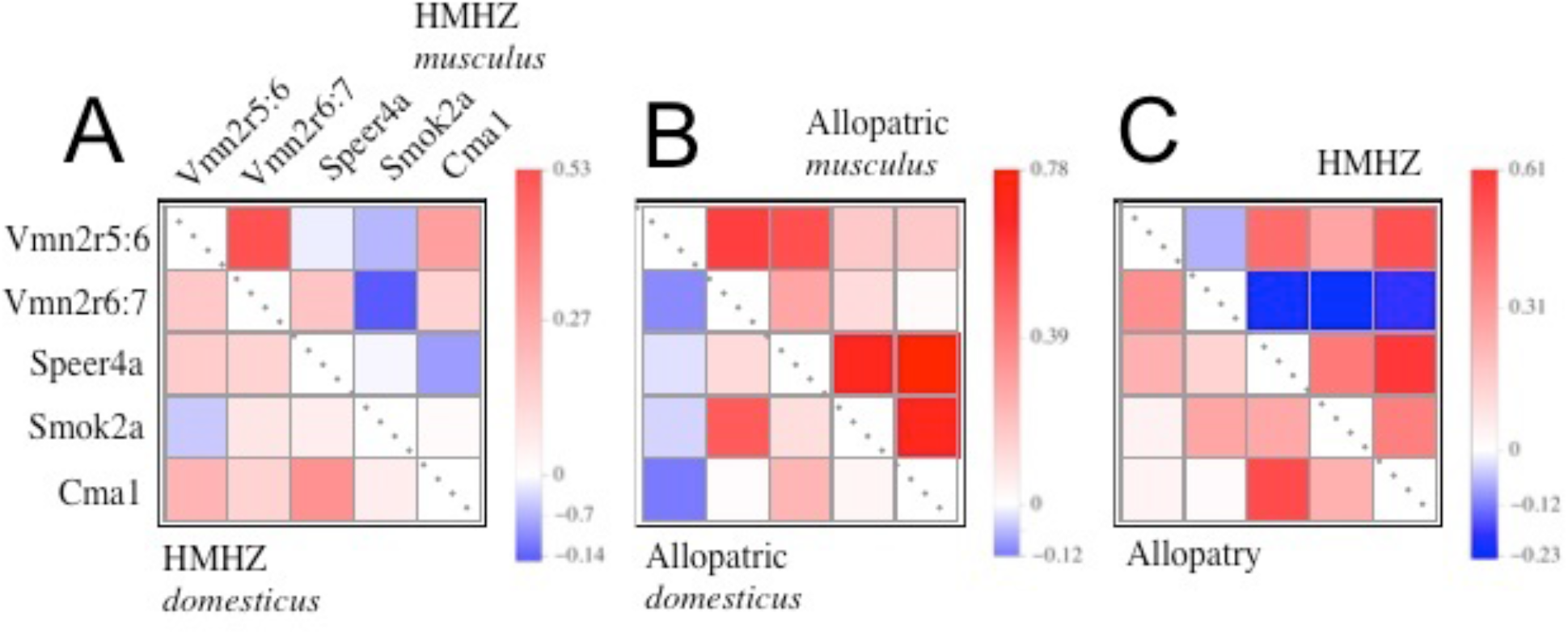
Matrix of pairwise covariances of gene copy numbers ( *D*_cov_) within different geographic sample groups. **A** – covariance within the sides of HMHZ (*musculus* and *domesticus*, above and below the main diagonal, respectively); **B** – covariance within the allopatric *musculus* and *domesticus* samples, above and below the diagonal, respectively; **C** – covariance within HMHZ as a whole (below diagonal) and all the allopatric samples grouped together (above diagonal). The order of gene names shown next to the matrix cells in **A** also applies to **B** and **C** panels.

We further explored the associations between CNV genotypes in respect to the variation in *HI*. Note that since only a few individuals in our study share the same sampling location (Baird et al., *in prep*.), we do not have a natural way of grouping the individual data to compare the center of the hybrid zone and its peripheries. Instead, we calculated *D*_cov_ cumulatively along the list of individuals sorted by the clinal variation. That is, the first value of *D*_cov_ was calculated for just 3 individuals with the highest (lowest) *HI* on one end of the list, the second for four and so on, thus progressing towards the other end of the list by including the next individual each time. The cumulative mean *D*_cov_ calculated from *musculus* to *domesticus* (left to right, blue line) and from *domesticus* to *musculus* (right to left, red line) is shown on Figure 2E: note a sudden increase in the mean *D*_cov_ between *Cma1, Speer4a, Smok2a* and *Vmn2r5:6* at the point of transition between the *musculus* and *domesticus* sides of HMHZ. The highest mean *D*_cov_ = 0.45 was observed when the entire *musculus* and *domesticus* sides were included in the calculation. All these genes have higher average copy numbers on the *musculus* side (Table 1). In contrast, the *Vmn2r6:7* gene is represented by a higher number of copies on the *domesticus* side and shows negative mean *D*_cov_, once both sides of HMHZ are included in the covariance estimator (mean *D*_cov_ = −0.19, Figure 2F).

#### Genome-wide association between SNP and CNV genotypes

Since neither the precise locations of all gene copies that amount to CNV nor the potential genome-wide epistatic interactions of the CNV genes are known, we searched for the possible links to CNV beyond the reference positions of the target gene. We examined the association of seven CNV genes with the inferred ancestral state of each of the ~500k SNPs across the entire genomes of 207 mice from the HMHZ (Baird et al. *in prep*.). For each SNP, the individuals were grouped according to their genotype (i.e. *musculus* homozygote [*mm*], heterozygote, [*md*], or *domesticus* homozygote [*dd*]), and a pairwise G-test (*mm-md*, *md-dd* and *mm-dd*) was applied to compare the distributions of copy numbers between groups (see Methods), based on the null hypothesis that both groups are sampled from the same distribution (Sokal and Rohlf 1995). Since the CN range at any one gene is the same for all SNPs, the absolute value of the G-test indicates how well a per-SNP genotype predicts the copy number state of a given CNV gene. The G-test is only bounded by zero, i.e. comparing two groups with identical distributions of CNs yields G = 0, while its upper value increases with the range of CNs that are being tested.

The G-test between the two homozygotes (*mm*-*dd*) was highly significant (Holm-Bonferroni adjusted p-value < 0.01) for 93-99% SNPs when compared to the CN data at five diagnostic genes. Except of Ifit3|3b and Defb7, the distributions of G-values were negatively skewed (i.e. most G-tests yielded higher values than the respective medians), with the modes ranging from G = 23 (*Vmn2r6:7*) and G = 65 (*speer4a*, Figure 4). Consistent with the fact that there is a large deficit of heterozygotes at >99% of all SNPs (Baird et al., *in prep*.), only 2 – 7 % of SNPs yielded significant G-tests in comparisons of the heterozygote with either homozygote (the tests *mm-md* and *md-dd*, Suppl Table 1). The high-copy-number *Speer4a* was a notable exception, with all three G-tests being significant for the vast majority of SNPs. Only 130 SNPs were significant in a G-test (*mm-dd*) on *Ifit3|3b* gene; 24 and 6 of these, respectively, were also significant in *mm-md* and *md-dd* comparisons. The *Defb7* gene yielded no significant result in any test.

**Figure 4.**
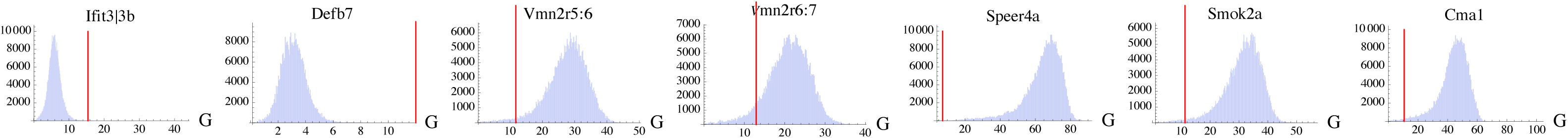
Distribution histograms of the [*mm-dd*] G-test (G) over the entire high-density SNP dataset (i.e. all 20 chromosomes pulled together), for each CNV gene. The red vertical line indicates the p = 0.01 significance threshold for each gene, after the Holm-Bonferroni correction. Note that (i) none of the G-test is significant for *Defb7* and only a small proportions of SNPs is significant for *Ifit3|3d;* (ii) distribution is negatively skewed for *Cma1, Smok2a, Speer4a, Vmn2r5:6* and *Vmn2r6:7*.

#### Introgression decreases association between the SNPs and gene copy number

The high-density SNP dataset allows for a fine-scale mapping of introgressed regions in the mouse genome (Baird et al. *in prep*). To increase the resolution of our analysis, we focused on the inrogression per chromosome rather than at the genome-wide level: a SNP allele was considered introgressed on its own chromosome if its source of origin (*musculus* or *domesticus*) contrasts with the majority of SNP alleles on that chromosome. For example, a *musculus*-derived allele on chromosome 4 carried by an individual with *HI*4 > 0.5 is considered introgressed on that chromosome. Note that the variation of marker genotypes across HMHZ is highly concordant for all autosomes (Macholán, Munclinger, Sugerková, et al. 2007; Baird et al., *in prep*.), and none of our CNV genes reference positions is sex-linked. Since the introgressed SNPs confuse the genome-wide distinction between *musculus* and *domesticus*, we expect them to be poor predictors of the copy number states of the diagnostic CNV genes. To verify this hypothesis, we designed a per-SNP additive estimator of introgression, *D*_int_, which accounts both for the total fraction of introgressed alleles at a given SNP as well as for the depth of introgression. Calculated over all 207 HMHZ individuals, the *D*_int_ demonstrates strong negative correlation with the G-test (*mm-dd*), on each chromosome and each of the five diagnostic genes (Spearman’s rank correlation coefficient < − 0.5, Suppl.Table 1). Similarly, the G-tests in two other SNP genotype pairs (*mm-md* and *md-dd*) were negatively correlated with *D*_int_ at *Speer4a* gene, but low number of significant G-tests (*mm-md* and *md-dd*) prevented the comparison at the other genes. We found no specific pattern of association that would distinguish any particular chromosome, including the X.

#### Outliers in the genome-wide SNP-CNV association map the reference position of Cma1 gene

We aimed to reveal those SNPs that demonstrate higher than expected association with the CNV data. Such SNPs can be identified as outliers in the genome-wide trend for a particular CNV gene. For that purpose, we first fitted a negative exponential model to the G-test and *D_int_* per-chromosome data for all SNPs per each diagnostic CNV gene (Figure 5, Suppl. Fig. 2). Those SNPs that appeared > 2 standard deviations from the fitted curve on the plot were shortlisted as preliminary outliers; a one-sided Grubbs’ test was then performed on the model residuals to detect the presence of at least one outlier in the dataset (Sokal and Rohlf 1995). The Grubbs test was applied repeatedly after removing the SNPs with the most extreme value one-by-one, until the test lost significance at the 99% confidence level.

**Figure 5.**
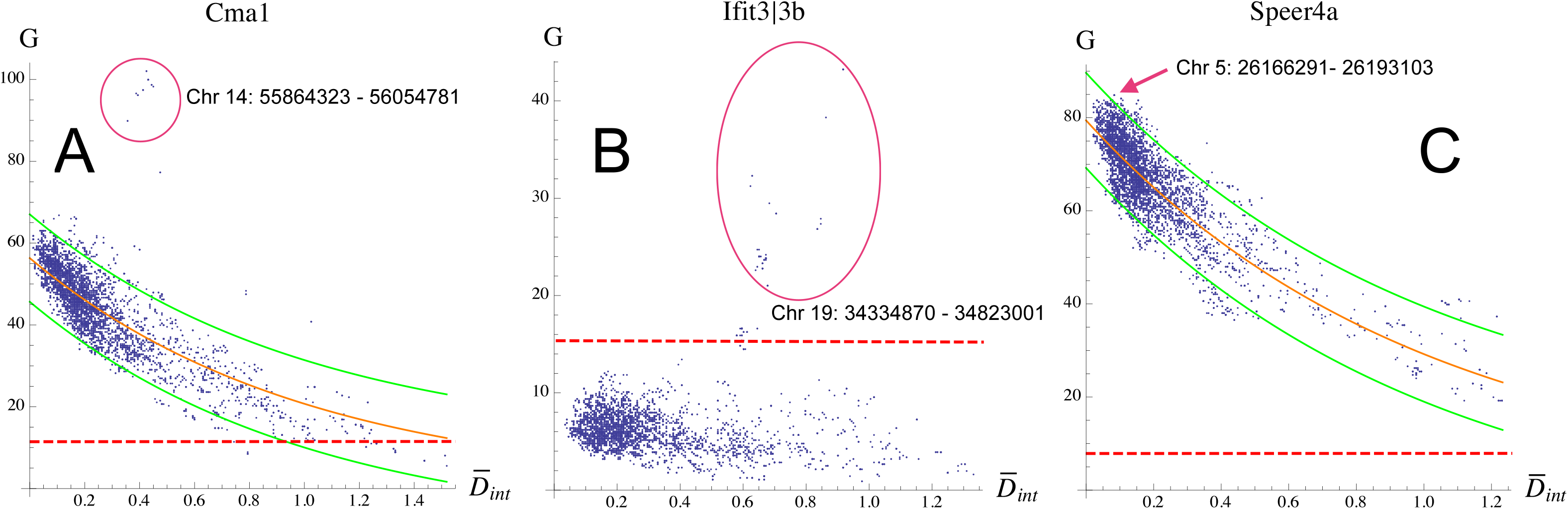
Chromosome-wide relationship between the per-SNP introgression index, *D*_int_, and the [*mm-dd*] G-test (G), in three cases where the reference positions of the CNV genes were mapped by the G-test maxima. The horizontal red line indicates the p = 0.01 significance threshold for each gene, after the Holm-Bonferroni correction, while the orange and green lines represent the negative exponential model and its 95% confidence intervals, respectively, fitted to the data. **A** – *Cma1* on Chromosome 14. Note that the region with the maximum G-test is also the outlier in the model fit; **B** – *Ifit3|3b* on Chromosome 19, **C** – *Speer4a* on Chromosome 5.

Without using any prior information on the reference positions of the CNV genes, we detected several continuous genomic regions mapped by the outlier SNPs (Table 2). The most notable group of outliers in a G-test with the copy number data of the *Cma1* gene is found on the Chromosome 14. This narrow region (~190 kb) shows moderate introgression from *musculus* to *domesticus (D_int_* ~ 0.4, Figure 5A, Figure 6A), and contains the precise annotated reference position of *Cma1*, along with 5 other reference genes (Table 2, Figure 6A). The region also possesses the highest genome-wide G-value among all CNV genes (G = 102.08). The distribution of gene copy numbers among the three diploid genotypes at the respective SNP revealed three distinct modes: CN ~ 2 for *domesticus* homozygotes, CN ~ 3 for heterozygotes and CN ~ 4 for *musculus* homozygote (Fig. 6D).

**Figure 6.**
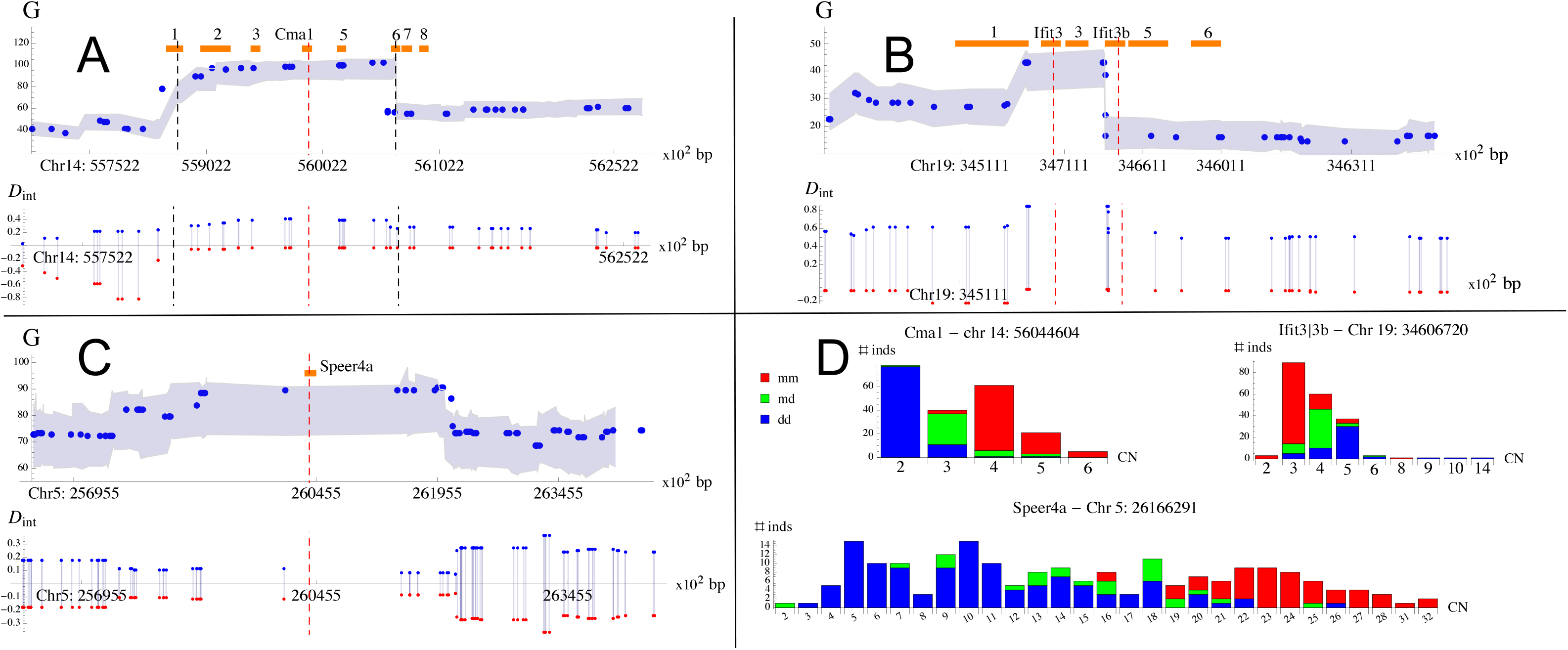
Detailed view of the genomic regions with reconfirmed CNV gene reference positions. **A** – Upper panel: per-SNP G-tests and *D*_int_ for *Cma1* on Chr 14. The blue dots represent distinct SNPs with their chromosomal positions shown on the x-axis. The corresponding G-test 95% confidence intervals are outlined by the shaded area. The region with the outlier G-tests (see Figure 5A, Table 2) is indicated by black vertical lines, while the red line marks the position of the amplicon used in ddPCR to genotype *Cma1* copy number. The annotated genes (Mouse Genome Database) in the region are shown by the orange boxes: 1 – *Cbln3*, 2 – *Khnyn*, 3 – *Sdr39u1*, 4 – *Cma1*, 5 – *Cma2*, 6 – *Mcpt1*, 7 – *Mcpt9*, 8 – *Mcpt2*. Lower panel: the per-SNP introgression index, *D*_int_, calculated separately on the *domesticus*and *musculus* sides (see Methods), shown by the blue and red dots, respectively. **B** – the same for Ifit3|3d gene on Chromosome 19, note that the *Ifit3|3b* ddPCR assay amplifies the target in both *Ifit3* and *Ifit3b* genes, hence the two red vertical lines. Numbered genes are: 1 – *Ifit2*, 2 – *Ifit3*, 3 – *Ifit1bl1*, 4 – *Ifit3b*, 5 – *Ifit1bl2*, 6 – *Ifit1*. **C** – the same for *Speer4a* gene on Chromosome 5. **D** – distributions of *mm* (red), *md* (green) and *dd* (blue) diploid genotypes among the copy numbers of *Cma1, Ifit3|3b* and *Speer4a*, at the SNP with the maximum G-test. The copy numbers on the x-axis were rounded to the closest integer.

**Table 2.** Candidate regions with stronger-than-expected association between CNV and SNP genotypes. See Supplementary Table 1 for reference genome position of the CNV loci

| position | CNV locus | G-test | *Dint* | Grubbs test (p < 0.01) | annotated genes in the region |
| --- | --- | --- | --- | --- | --- |
| **chr14:** 55864323 - 56054781 | ***Cma1*** | 77.47-102.08* | 0.353-0.474 | 12.037 | ***Cma1***, *Cma2*, + more |
| chr6: 21409624-21416010 | *Cma1* | 62.8 | 0.526 | 6.001 | *Kcnd2* |
| chr2: 42875644- 43568817 | *Smok2a* | 49.2-56.1** | 0.432-0.547 | 7.034 | *Gm30225(ncRNA)* |
| chr8: 79530293- 79666745 | *Smok2a* | 35.53, 42.55 | 0.356, 0.743 | 5.422 | *Gm39228*, *Gm8077*, *Gm8079* |
| chr11: 117665934-117754065 | *Vmn2r6* | 28.01-32.01 | 0.368-0.475 | 5.205 | *Tnrc6c* |
| chr15: 53961638-54045364 | *Vmn2r6* | 22.93, 26.67 | 0.695-0.9 | 5.876 | *Gm26933* |
| **chr19:** 34334870 - 34823001 | ***Ifit3\|3b*** | 14.56-43.23* | 0.582-0.916 | n/a | ***Ifit3***, ***Ifit3b*** + more |
| chr4: 134357389 | *Ifit3\|3b* | 15.53 | 0.758 | n/a | *Slc30a2* |
| chr8: 106855553- 106895716 | *Ifit3\|3b* | 15.69-16.15 | 0.327-0.336 | n/a | *Tango6* |
| chr12: 36324689- 36653266 | *Ifit3\|3b* | 14.68-15.74 | 0.234-0.303 | n/a | *Ispd* + 5 more |
| **chr5:** 26201557- 26203039 | ***Speer4a*** | 90.41* | 0.137 | n/a | ***Speer4a*** (1.56 kb upstream) |

\* genome-wide maximum for that locus \*\* maximum on that chromosome

#### Reference positions of Speer4a, Ifit3 and Ifit3d genes mapped by the G-test maxima

In cases where the outlier analysis was not significant or could not be performed, we examined closely the genomic regions that contained the SNPs with the maximum genomewide G-values for each CNV gene. This simple approach revealed a small continuous region on Chromosome 5 with the maximum of G = 90.41 (*mm-dd* test with *Speer4a* CN data, Figure 5C) displayed by a SNP located only 1.56 kb away from the reference position of the *Speer4a* gene (Figure 6C). At this SNP, several modes are observed in the distribution of gene copy numbers for each SNP genotype, however, the largest CN mode in heterozygotes (CN ~ 18) was intermediate in respect to the homozygotes (largest modes in *dd* CN ~ 5 and CN ~ 9, largest mode in *mm* CN ~ 23, Fig. 6D).

Similarly, the genome-wide maximum G-value for the *Ifit3|3b* CN data was observed close to the reference locations of the *Ifit3* and *Ifit3b* genes (Figure 5B, Figure 6B). In a G-test with the *Ifit3|3d* CN data, the two neighboring SNPs with the genome-wide maximum G = 43 are located in the 3’ and 5’ flanks of the reference position of *Ifit3*, respectively, and in the 3’ flank of the *Ifit3b* (Table 2, Figure 6B). Notably, both are located in a region of deep introgression from *musculus* to *domesticus* (*D_int_* ~ 0.8, Figure 5B, Figure 6B). The distribution of the copy numbers at the two focal SNPs had the following modes: CN ~ 5 for *dd* homozygotes, CN ~ 4 for *md* heterozygotes and CN ~ 3 for *mm* homozygotes (Figure 6D). Recall that (i) our *Ifit3|3b* primer assay has equal affinity to both *Ifit3* and *Ifit3b* reference genes (Suppl Table 1), and (ii) we found no association between the *Ifit3|3d* CN data and the hybrid index. Since only a small number of other SNPs were significant in the G-test with *Ifit3|3b* copy number data, we characterized their genome locations as well (Table 2).

#### Outlier SNPs point to multiple regions beyond the reference positions of the CNV genes

A number of other regions contained the outlier SNPs in the G-tests with *Cma1, Smok2a* and *Vmn2r6:7* CN data, with no apparent links to the reference positions of the respective genes (Table 2). In all these cases, the outlier region resided on a different chromosome than the corresponding location of the CNV gene in the reference. One outlier region on Chromosome 2 also had the genome-wide maximum G-value in a test on the *Smok2a* CN data: note that the reference position of the *Smok2a* gene is on Chromosome 17. The complete list of the outlier regions is given in Table 2.

## Discussion

We described distinct patterns of gene copy number variation of several protein-coding genes between *M. m. musculus* and *M. m. domesticus*, both in allopatry and across the contact zone in Central Europe. To our knowledge, this is the first survey of autosomal gene CNV in any hybrid zone: although we only examined a small set of gene markers, our results support the current view on CNV as a major component of genetic structure within and between populations (Scavetta and Tautz 2010; Schrider and Hahn 2010; Iskow, Gokcumen, and Lee 2012; Bryk and Tautz 2014; Redon et al. 2006; Chain et al. 2014). While no common CNV pattern was detected in all markers, five genes showed prominent CNV structure between the sides of the hybrid zone, which in three cases extended to the allopatric populations. By examining associations between the CNV and SNP genotypes across the hybrid zone, we were able to confirm the precise locations of three CNV markers, as annotated in the reference house mouse genome. The same approach revealed several genomic locations with highly significant associations but previously not reported to have any relationship to the respective CNV genes.

### The candidate gene approach led to a high CNV discovery rate

We were able to detect CNV in 7 out of 11 candidate genes tested initially (see Methods) in a single transect across HMHZ. Such a high success rate of CNV discovery may reflect: (i) very high genome-wide level of CNV between *M.m. musculus* and *M.m. domesticus* and/or (ii) prevalence of inter-population CNV among the candidate genes chosen for copy number typing. Previously, the average reported proportion of the house mouse genome affected by CNV was ranged from ~ 2.3 % (Henrichsen et al. 2009) to 6.87% (Locke et al. 2015). Note that all genome-wide CNV surveys up to date dealt with small samples of wild-caught mice, and only represented *Mus. m. domesticus* (Pezer et al. 2015; Locke et al. 2015; Henrichsen et al. 2009; Keane et al. 2011). Since the copy number differences tend to accumulate with divergence (Gokcumen et al. 2011; Chain et al. 2014), we would expect higher CNV between the house mouse subspecies than within any one of them. However, it is equally likely that the high CNV discovery rate in our study is mostly due to the candidate gene approach (ii), i.e. sampling the genes from the previously reported mouse CNVRs. An important question is what part of this success is owed to our arbitrary focus on the putative speciation genes. Since we don’t have a reliable control group of “speciation-irrelevant” genes, it can’t be answered conclusively, nevertheless, our results are in line with previously suggested importance of CNV at the early stages of speciation (Chain et al. 2014).

### The vomeronasal receptors may be under reinforcement selection in the HMHZ

Among the seven genes that displayed CNV in the set of 252 individuals, five were found to be strongly associated with genome contributions of *M. m. musculus* and *M. m. domesticus* in the zone of contact: for three genes (*Cma1, Smok2a* and *Speer4a*), the same patterns also extended to the distant allopatric samples from both *M. m. musculus* and *M. m. domesticus* geographic ranges. Intriguingly, two CNV markers, *Vmn2r5:6* and *Vmn2r6:7*, despite having clear copy number differences between the sides of the hybrid zone, do not differentiate between the allopatric samples of the respective “pure” subspecies. This observation is consistent with the scenario of reproductive character displacement, found in many contact zones (Pfennig and Pfennig 2009; C. Smadja and Ganem 2005) and potentially caused by reinforcement selection favoring stronger prezygotic isolation between the hybridizing taxa (Servedio and Noor 2003; Bímová et al. 2011). An alternative mechanism for the character displacement, e.g. resource competition between the distinct phenotypes, cannot be ruled out in this and other similar cases: however, the role of vomeronasal receptors in pheromone reception and mate recognition provides support the reinforcement scenario in the house mouse (Del Punta et al. 2002). In the agreement with our observations, no significant differences of *Vmn2r5* and *Vmn2r6* copy numbers was found between small samples from France, Germany and Iran (Pezer et al. 2015). Smadja et al. (2015) compared the levels of microsatellite allele diversity in genomic regions containing the MUP (Murine Urinary Proteins) and VR (vomeronasal receptors) gene families, between the natural populations of the house mouse sampled at various distances from the hybrid zone in Denmark. The authors found that several genes from the VR2 subfamily displayed particularly low allelic diversity in the hybrid zone, as opposed to the distant samples, which could be seen as an evidence for a hard sweep caused by the reinforcing selection. Note that the genes *Vmn2r5, 6* and *7* examined in our study also belong to the VR2 subfamily, but don’t share the same chromosome with any of the VR2 genes detected by Smadja et al. (2015).

### Possible evidence for introgression of Cma1 and Ifit3|3b CN haplotypes from musculus to domesticus

Unlike the vast majority of SNPs located in the introgressed regions, those mapping the reference position of the *Cma1* gene on Chromosome 14 show very high association with the copy number of the same gene. The CN distribution among the three SNP genotypes is consistent with one extra copy of the gene hypothetically located very close to the reference position and originated from *M. m. musculus* (*Cma1* 2n = 4), while *M. m. domesticus* would possess just one copy (*Cma1* 2n = 2). Note that individual copy numbers of *Cma1* in our ddPCR assays ranged from 1 to 6 (Table 1), however these CN types occurred at much lower frequencies. Most importantly, the haplotype containing two linked copies of *Cma1* seems to introgress into *domesticus* genome as marked by the SNP genotypes at this position. This claim is indirectly supported by the observation that *Cma1* diploid copy numbers almost invariably equal two and four in the respective *domesticus* and *musculus* allopatric populations that we sampled. At the same time, mice genotyped by Pezer et al. (2015) in France all had four or (rarely) three diploid copies of *Cma1*, while all individuals from Germany and Iran had two copies. Such geographic pattern may have been caused by (i) ancestral CNV preceding *musculus*-*domesticus* split, (ii) long range introgression and (iii) *de novo* duplication of *Cma1* in France. Both the strong population structure in Western Europe observed by Pezer et al. (2015) and the introgression pattern among linked SNPs at HMHZ (Baird et al. *in prep*.) make ancestral polymorphism (i) an unlikely explanation: differentiating between the other two scenarios (ii-iii) is not possible with the current data.

The gene pair *Ifit3* and *Ifit3b* represents another case of strong association between the SNP and the copy number data, suggesting that a CNV haplotype consisting of closely linked gene copies exists in this reference genome region. Recall that the same region is characterized by massive introgression from *musculus* to *domesticus*. Since the SNP genotypes here do not correlate with the hybrid index, neither do the *Ifit3|3b* copy number data. This explains why we did not observe any copy number differences at *Ifit3|3b* between the sides of the HMHZ. It is tempting to speculate that the two ancestral populations of the house mouse differed in the *Ifit3|3b* copy number before secondary contact, but since then the higher CN haplotype on the *domesticus* side has been replaced by the lower CN haplotype derived via introgression from the *musculus* side, eroding the association signature everywhere but within the inrogressed region. To consider this scenario one would have to assume that the hypothetical ancestral populations were distinct in respect to *Ifit3|3b* from the present-day allopatric populations, since we also did not detect any significant inter-population differences in *Ifit3|3b* CNs outside HMHZ, nor did Pezer et al. (2015), who only genotyped the *Ifit3* locus. An alternative scenario involving *de novo* origin of the lower CN haplotype on the *musculus* side with subsequent introgression cannot be excluded. Note that the CN distribution at the *Ifit3|3b* gene is characterized by a long right tale (up to 14 copies in one individual, Figure 6D), which might suggest that recurrent CNV mutations are relatively common in these genes.

### Outlier analysis may point to previously unknown locations of gene copies

In this pilot study, we confirmed the known reference and found several new genomic locations with possible involvement in CNV (Table 2). Our approach differed from that used in the conventional genome-wide association studies (GWAS), which usually examine the associations between the phenotype and a particular SNP allele. Instead, we focused on the *derived ancestral source* of the SNP (i.e. *musculus* or *domesticus* ancestral population, Baird et al. *in prep*) and identified the outliers in the genome-wide negative correlation trend between the diagnostic gene copy number and introgression. Out method is thus suited for detecting the signal of a specific scenario of population admixture, where (i) the ancestral populations differ in gene copy number and (ii) the subsequent contact leads to introgression of CN haplotypes. Indeed, the *Cma1* and *Ifit3|3b* genes can be regarded, albeit speculatively, as examples of such a scenario developing at different stages: in case of *Cma1*, the introgression might be ongoing but the population differences in copy number are still maintained, whereas the CN differences at *Ifit3|3b* have been already eroded, leaving only the association signature within the region of introgression. The new genomic regions mapped by the outlier SNP could potentially harbor additional gene copies, or be involved in CNV-specific epistatic interactions. Two other observations call for caution: first, unlike at the confirmed reference locations, the outlier G-tests at the new regions are below the respective genome-wide maxima, and second, none of the new regions share the same chromosome with a reference location of respective CNV gene. The latter observation is particularly interesting since it seems to preclude the most frequent mechanisms of CNV origin by gene duplication, i.e. non-homologous recombination (NAHR, Hastings et al. 2009). Occurrence of nearly identical gene copies on different chromosomes can be explained by transposition, although this mechanism is less common (Kaessmann, Vinckenbosch, and Long 2009). We suggest that sequencing and *de novo* assembly of the newly discovered CNV-associated regions, in a small sample of individuals with confirmed difference in the copy number, may resolve whether the outlier signals are caused by the putative extra gene copies. Investigation in that direction is currently underway.

## Material and Methods

### Primer design and initial CN genotyping

Our ability to design specific primers each with at least 3 basepair mismatches to the next most likely target amplicon <1kb length was one of the major criteria in choosing the candidate genes for CN genotyping. Primer and fluorescent probe design was performed using the Primer3 software (http://bioinfo.ut.ee/primer3-0.4.0)and their specificity verified by NCBI’s Primer-BLAST tool (http://www.ncbi.nlm.nih.gov/tools/primer-blast) against the mouse genome reference assembly mm10 (organism selected as *Mus*, taxid:10088). Many members of large gene families did not satisfy the amplification specificity criterion (iv), further narrowing the list down to a few tens of candidate genes (list is available upon request). Primers targeting the coding regions or the flanks of 11 genes (Suppl.Table 1) were initially tested in singleplex ddPCR reactions using 12 randomly chosen individuals from HMHZ, and one individual from each of the C3Hb, C57BL/6J (both classical laboratory strains), SMON and XBS strains (derived from wild *M. spretus* and *M. macedonicus*). Among these, the *Plac9a, Tex35, Spaca4* and *Defb8* genes appeared to have uniformly similar copy numbers and were therefore not selected for further genotyping. The remaining seven amplicons, targeting the *Cma1, Smok2a, Speer4a, Ifit3, Ifit3b* and *Defb7*, as well as two reference locations between *Vmn2r5:6* and *Vmn2r6:7* genes showed copy number differences in the initial analysis and so were genotyped for the entire 252 sample set.

### Droplet Digital PCR

The ddPCR reactions and the initial PCR optimization experiments were performed on the QX100/200 Droplet Digital PCR System (Bio-Rad, Hercules, CA, USA). The copy numbers of all CNV genes in our study were estimated relative to a fragment of the *Tert* gene, which is known to be present in two copies per diploid genome (Pezer et al. 2015). All CNV assays, as well as restriction endonucleases in each reaction, were chosen to be compatible with the *Tert* assay (Bio-Rad 2015). Two techniques were used, singleplex and multiplex, and the data obtained were directly compared (results are available upon request). While the degree of correspondence between the two methods varied across amplicons, in each case the singleplex method allowed for reliable detection of the CNV among samples. For singleplex ddPCR reactions, 60 ng of genomic DNA template were added to the master mix of 20 μl containing the Bio-Rad 2× ddPCR™ Supermix QX200™ EvaGreen^®^ and the primers (SIGMA_®_,St. Louis, MO, USA) at the final concentration of 450-900 nM. To maximize the accuracy of resolving the copy number in singleplex, we attempted to equalize all reagent concentrations within a pair of reaction mixtures, one mixture containing the primers for the target amplicon and the other containing the primers for the reference *Tert* gene. The reaction mixtures were then converted to droplets: to minimize the potential bias at this step, we ensured that the reference/target pair is not split between different runs of the QX100/200 ddPCR Droplet Generator machine. The PCR was performed on the resulting pairs of droplet mixtures, using the following protocol: 5 min at 95°C (1 cycle), 30 sec at 95°C + 1 min at 57-60°C (40 cycles), 5 min at 4°C (1 cycle) and 5 min at 90°C (1cycle). The absolute number of droplets with positive amplification signal was estimated separately for the target and the reference on a QX200 Droplet Reader, using the Bio-Rad’s Quantasoft™ Software (version 1.6.6). The ratio between the absolute concentrations of the target and the reference was then used to infer the approximate genomic copy number of the target. Later, the data obtained in singleplex were directly compared to those from the much more precise multiplex ddPCR reactions (results are available upon request). While the degree of correspondence between the two methods varied across amplicons, in each case the singleplex method allowed for reliable detection of the CNV among samples.

For the multiplex ddPCR we used the same primer sequences as in singleplex. The primers for the multiplex were ordered separately as a part of the Prime Time®Std qPCR Assays (Coralville, Iowa, USA) that also included a fluorescent hybridization probe. The probes for the target amplicons were labeled with FAM, and the probe for the reference *Tert* gene was labeled with HEX. We used 15-100 ng of genomic DNA template in 20 μl reactions containing the 2× ddPCR™ Super Mix for Probes. The final concentrations of the primers and the probes were 450-900 nM and 125-250 nM, respectively. After converting the multiplex reaction mixtures into droplets, the amplification was performed using the following protocol: 10 min at 95°C (1 cycle), 30 sec at 94°C + 1 min at 57 – 60°C (40 cycles) and 98°C for 10 min (1 cycle), with a ramp speed of 2.5°C/sec. Each amplicon was characterized by a unique and consistent distribution of the fluorescent signal frequency (Suppl.Fig. 1). This copy number genotypes and the measurement errors were then resolved on the QX200 Droplet Reader using the corresponding option in the Quantasoft™ Software.

#### Digestion of DNA templates

To ensure the independent separation the genomic copies of the CNV loci among droplets, the DNA templates were digested directly in the ddPCR master mixture, as per the recommendation of Bio-Rad (Bio-Rad 2015): we used 0.1 unit of BamHI or MseI restriction endonucleases (New England Biolabs®, Hitchin, Hertfordshire, UK) per reaction, and incubated the master mixtures at 37**°**C for 10 min as the last step before converting the mixtures into droplets. Separate experiment was performed on eight DNA samples randomly chosen form the HMHZ sample set to compare the effectiveness of the copy number resolution of the locus *Smok2a* after the in-mixture digestion relative to the ddPCR on the pre-digested template. The pre-digestion was performed for one hour at a concentration of 100 ng of DNA per 50-μl reaction, using 5 units of MseI enzyme per 1 μg of DNA and 5 μl of 1X SmartCut™ Buffer (New England Biolabs®). This was followed by the incubation for 45 min at 37°C and inactivation for 20 min at 65° C. The pre-digested template was then subjected to the same multiplex ddPCR protocol using the *Smok2a* primer assay, but without adding the restriction enzyme to the mixture and skipping the additional 10 min incubation step. After the completion of ddPCR, neither the copy number estimates nor the corresponding measurement errors were significantly different between the two experiments (Pearson chi-square test *p* = 0.56, Suppl.Fig. 2).

### The 252 high-density MDA-SNP sample set

We genotyped 43 individuals from the allopatric populations of *Mus m. musculus* and *Mus m. domesticus*, sampled across Europe and Middle East, one individual of *M. spicilegus* from Serbia and one individual of *M. macedonicus* from Lebanon, and 207 individuals from the Czech-Bavarian transect of HMHZ (Figure 1). The genetic structure of this transect has been investigated previously (Macholán et al. 2007, 2008, 2011; Dufková et al. 2011; Janousek et al. 2012, 2016), Baird et al. 2016) analyzed the contributions of inferred *musculus* and *domesticus* source populations in the genomes of the same 252 individuals, using a high-density MDA (Mouse Diversity Array) with approximately 500k SNPs (see Baird et al. in prep. for details of the DNA extraction, MDA-SNP chip analysis procedures and the method of inference). The data used here (referred to as 252 SNP dataset in the following) includes the source label (*musculus* or *domesticus*) per each SNP allele, and the chromosomal position of each SNP. Each individual in the 252 SNP dataset is thus characterized by the hybrid index (*H_t_*), calculated as the proportion of diagnostic SNP alleles sourced from the *domesticus* population (i.e. a genome carrying 100%*musculus* alleles has *H_t_* = 0 and that with 100%*domesticus* alleles SNPs has *H_t_* = 1, the estimate is based on ~500k SNPs). The hybrid index *Hc* is similarly calculated for each chromosome (Baird et al. *in prep*.). A vast majority of diagnostic SNPs are thought to have diverged since the split between *M.m. musculus* and *M.m. domesticus* ~ 0.5 myr ago (Geraldes et al. 2008), while a tiny proportion may represent ancestral polymorphism (Baird et al. *in prep*.).

### Inference of population structure

#### Relative distance between populations

The pairwise per-locus *V_ST_* statistic (Redon et al. 2006) was calculated as 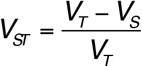, where *V_ST_* is the variance in the estimated copy numbers among all individuals in a given dataset, and 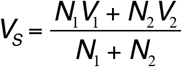 is the average variance within each population (*V*), weighted by the sample size (*N*).

#### Covariance between loci

The normalized pairwise covariance *D*_cov_ between the two sets of copy number estimates (*loc*) was calculated as 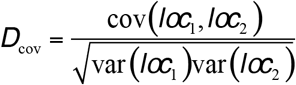.

#### The per-sNP G-test

We used a pairwise G-test as a measure of independence of the copy number distributions between two SNP genotypes, calculated as:

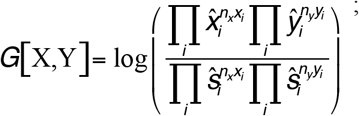

where **X** and **Y** are the two vectors of the same length, representing the numbers (*x,y*) of individual copy number genotypes that fall in the arbitrarily defined bins of length = 1, ranging from the CN = 0.5 to the maximum CN for a given locus. An individual CN estimate is thus effectively rounded to the nearest whole number. The “hat” symbol represents the values of the respective vector normalized between 0 and 1. Our expectation for the CN distribution is calculated as 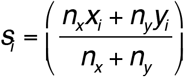, where 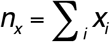 and 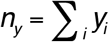 are the respective sums of rounded copy numbers. Since ***G*** is a pairwise test, we made three comparisons: between two SNP homozygotes (*mm* and *dd*) and between the heterozygote (*md*) and either *mm* or *dd*. The latter of these, however, are much less informative due to the overall low numbers of heterozygotes in our dataset, and were therefore not included in the final analysis.

#### Estimating the degree of introgression on a given chromosome and fitting the negative exponential model to G-test

We calculated the per-SNP introgression index, *D_int_* as follows: 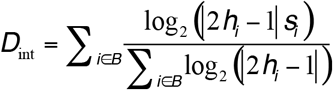; where *h_i_* is a chromosomal hybrid index of the *i*-th individual and ***S_i_*** is a number of introgressed alleles (i.e. 0, 1 or 2) carried by that individual at a given SNP. To draw the introgression diagrams on Figure 6, two estimates, 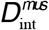 and 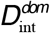 on the *musculus* and *domesticus* sides of HMHZ, respectively, were calculated separately, i.e. for two groups of individuals with *h_i_* < ½ and *h_i_* > ½. The two estimates were averaged (*D*_int_) in order to fit the negative exponential model 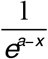, where *a* is a fitted parameter and *x* is the variable, to the *D*_int_/ G-test data (Figure 5).

## Authors contribution

AY designed the study, and performed statistical analysis with the help from SJEB. AY and ZH performed the laboratory analysis, and ZP contributed to primer design and development of ddPCR protocols. MM and JP provided DNA samples. AY wrote the paper, and all authors contributed significantly to the final version of the manuscript.

## Acknowledgements

We would like to thank Dr. Diethard Tautz for hosting AY and ZH at Max-Planck Institute for Evolutionary Biology (MPI), where the ddPCR experiments were conducted, and his helpful comments at every stage of the work. Joelle Göuy de Bellocq and Anna Bryja advised on the primer design. We especially grateful to Nicole Thomsen for her technical assistance at MPI. The ddPCR analysis was funded by the Max Planck Society. AY, SJEB and ZH were supported by the NextGenProject CZ [grant number 1.07/2.3.00/20.0303]. High-throughput computational capacity was provided by the Czech National Grid Infrastructure MetaCentrum running under the program “Projects of Large Research, Development, and Innovations Infrastructures” [grant number CESNET LM2015042]. The funding bodies played no role in determining study design, data collection, analysis, and interpretation, or in writing the manuscript.

## Literature Cited

Albrechtová, Jana, Tomás Albrecht, Stuart J E Baird, Milos Macholán, Geir Rudolfsen, Pavel Munclinger, Priscilla K Tucker, and Jaroslav Piálek. 2012. “Sperm-Related Phenotypes Implicated in Both Maintenance and Breakdown of a Natural Species Barrier in the House Mouse.” Proceedings. Biological Sciences / The Royal Society 279 (1748): 4803–10. doi:10.1098/rspb.2012.1802.

Baird, S. J. E. 1995. “A SIMULATION STUDY OF MULTILOCUS CLINES.” Evolution 49 (6). Wiley: 1038–45. http://cat.inist.fr/?aModele=afficheN&cpsidt=10891648.

Barton, N. H. 1983. “Multilocus Clines.” Evolution 37 (3): 454. doi:10.2307/2408260.

Barton, N. H., and G. M. Hewitt. 2003. “Analysis of Hybrid Zones.” http://dx.doi.org/10.1146/annurev.es.16.110185.000553. Annual Reviews 4139 El Camino Way, PO. Box 10139, Palo Alto, CA 94303-0139, USA.

Barton, Nick, and Bengt Olle Bengtsson. 1986. “The Barrier to Genetic Exchange between Hybridising Populations.” Heredity 57 (3): 357–76. doi:10.1038/hdy.1986.135.

Vošlajerová Bímová, Barbora, Miloš Macholán, Stuart J E Baird, Pavel Munclinger, Petra Dufková, Christina M. Laukaitis, Robert C. Karn, Kenneth Luzynski, Priscilla K. Tucker, and Jaroslav Piálek. 2011. “Reinforcement Selection Acting on the European House Mouse Hybrid Zone.” Molecular Ecology 20: 2403–24. doi:10.1111/j.1365-294X.2011.05106.x.

Bio-Rad. 2015. Droplet Digital PCR Application Guide. http://www.bio-rad.com/webroot/web/pdf/lsr/literature/Bulletin_6407.pdf.

Boursot, P. and K. Belkhir. 2006. Mouse SNPs for evolutionary biology: Beware of ascertainment biases. Genome Research 16:1191–1192.

Bryk, Jarosław, Mehmet Somel, Anna Lorenc, and Meike Teschke. 2013. “Early Gene Expression Divergence between Allopatric Populations of the House Mouse (Mus Musculus Domesticus).” Ecology and Evolution 3 (3): 558–68. doi:10.1002/ece3.447.

Bryk, Jarosław, and Diethard Tautz. 2014. “Copy Number Variants and Selective Sweeps in Natural Populations of the House Mouse (Mus Musculus Domesticus).” Frontiers in Genetics 5 (June): 153. doi:10.3389/fgene.2014.00153.

Cardoso-Moreira, Margarida, J. Roman Arguello, Srikanth Gottipati, L.G. Harshman, Jennifer K. Grenier, and Andrew G. Clark. 2016. “Evidence for the Fixation of Gene Duplications by Positive Selection in *Drosophila*.” Genome Research 26 (6): 787–98. doi:10.1101/gr.199323.115.

Chain, Frédéric J. J., Philine G. D. Feulner, Mahesh Panchal, Christophe Eizaguirre, Irene E. Samonte, Martin Kalbe, Tobias L. Lenz, et al. 2014. “Extensive Copy-Number Variation of Young Genes across Stickleback Populations.” PLoS Genetics 10 (12): e1004830. doi:10.1371/journal.pgen.1004830.

Conrad, Donald F, Dalila Pinto, Richard Redon, Lars Feuk, Omer Gokcumen, Yujun Zhang, Jan Aerts, et al. 2010. “Origins and Functional Impact of Copy Number Variation in the Human Genome.” Nature 464 (7289). Macmillan Publishers Limited. All rights reserved: 704–12. doi:10.1038/nature08516.

Cutler, Gene, Lisa A Marshall, Ni Chin, Helene Baribault, and Paul D Kassner. 2007. “Significant Gene Content Variation Characterizes the Genomes of Inbred Mouse Strains.” Genome Research 17 (12): 1743–54. doi:10.1101/gr.6754607.

Dean, MD, and MW Nachman. 2009. “Faster Fertilization Rate in Conspecific versus Heterospecific Matings in House Mice.” Evolution. http://onlinelibrary.wiley.com/doi/10.1111/j.1558-5646.2008.00499.x/full.

Del Punta, Karina, Trese Leinders-Zufall, Ivan Rodriguez, David Jukam, Charles J. Wysocki, Sonoko Ogawa, Frank Zufall, and Peter Mombaerts. 2002. “Deficient Pheromone Responses in Mice Lacking a Cluster of Vomeronasal Receptor Genes.” Nature 419 (6902). Nature Publishing Group: 70–74. doi:10.1038/nature00955.

Dod, Barbara, Carole Smadja, Robert C. Karn, and Pierre Boursot. 2005. “Testing for Selection on the Androgen-Binding Protein in the Danish Mouse Hybrid Zone.” Biological Journal of the Linnean Society 84 (3): 447–59. doi:10.1111/j.1095-8312.2005.00446.x.

Dzur-Gejdosova, Maria, Petr Simecek, Sona Gregorova, Tanmoy Bhattacharyya, and Jiri Forejt. 2012. “Dissecting the Genetic Architecture of F1 Hybrid Sterility in House Mice.” Evolution; International Journal of Organic Evolution 66 (11): 3321–35. doi:10.1111/j.1558-5646.2012.01684.x.

Emes, R.D., M. C. Riley, C. M. Laukaitis, L. Goodstadt, R. C. Karn, and C. P. Ponting. 2004. Comparative evolutionary genomics of androgen-binding protein genes. Genome Research 14 (8):1516–1529.

Eppig, J. T., J. A. Blake, C. J. Bult, J. A. Kadin, J. E. Richardson, and The Mouse Genome Database Group. 2015. “The Mouse Genome Database (MGD): Facilitating Mouse as a Model for Human Biology and Disease.” Nucleic Acids Research 43 (D1). Oxford University Press: D726–36. doi:10.1093/nar/gku967.

Firman, Renée C., Leigh W. Simmons, WR Rice, DJ Hosken, TW Garner, PI Ward, RC Firman, et al. 2014. “No Evidence of Conpopulation Sperm Precedence between Allopatric Populations of House Mice.” Edited by William J. Etges. PLoS ONE 9(10). Public Library of Science: e107472. doi:10.1371/journal.pone.0107472.

Forejt, Jiří. 1996. “Hybrid Sterility in the Mouse.” Trends in Genetics 12 (10): 412–17. doi:10.1016/0168-9525(96)10040-8.

Gay, Laurène, P. a. Crochet, Douglas a. Bell, and Thomas Lenormand. 2008. “Comparing Clines on Molecular and Phenotypic Traits in Hybrid Zones: A Window on Tension Zone Models.” Evolution 62: 2789–2806. doi:10.1111/j.1558-5646.2008.00491.x.

Geraldes, Armando, Patrick Basset, Barbara Gibson, Kimberly L Smith, Bettina Harr, Hon-Tsen Yu, Nina Bulatova, Yaron Ziv, and Michael W Nachman. 2008. “Inferring the History of Speciation in House Mice from Autosomal, X-Linked, Y-Linked and Mitochondrial Genes.” Molecular Ecology 17 (24): 5349–63. doi:10.1111/j.1365-294X.2008.04005.x.

Gokcumen, Omer, Paul L Babb, Rebecca C Iskow, Qihui Zhu, Xinghua Shi, Ryan E Mills, Iuliana Ionita-Laza, et al. 2011. “Refinement of Primate Copy Number Variation Hotspots Identifies Candidate Genomic Regions Evolving under Positive Selection.” Genome Biology 12 (5): R52. doi:10.1186/gb-2011-12-5-r52.

Hardouin, Emilie A, Annie Orth, Meike Teschke, Jamshid Darvish, Diethard Tautz, and François Bonhomme. 2015. “Eurasian House Mouse (Mus Musculus L.) Differentiation at Microsatellite Loci Identifies the Iranian Plateau as a Phylogeographic Hotspot.” BMC Evolutionary Biology 15 (1). BioMed Central: 26. doi:10.1186/s12862-015-0306-4.

Harr, B. 2006. “Genomic Islands of Differentiation between House Mouse Subspecies.” Genome Research 16 (6): 730–37. doi:10.1101/gr.5045006.

Hastings, P J., James R. Lupski, Susan M. Rosenberg, and Grzegorz Ira. 2009. “Mechanisms of Change in Gene Copy Number.” Nature Reviews Genetics 10 (8). Nature Publishing Group: 551–64. doi:10.1038/nrg2593.

Henrichsen, Charlotte N, Nicolas Vinckenbosch, Sebastian Zöllner, Evelyne Chaignat, Sylvain Pradervand, Frédéric Schütz, Manuel Ruedi, Henrik Kaessmann, and Alexandre Reymond. 2009. “Segmental Copy Number Variation Shapes Tissue Transcriptomes.” Nature Genetics 41 (4): 424–29. doi:10.1038/ng.345.

Iskow, Rebecca C., Omer Gokcumen, and Charles Lee. 2012. “Exploring the Role of Copy Number Variants in Human Adaptation.” Trends in Genetics 28 (6): 245–57. doi:10.1016/j.tig.2012.03.002.

Janousek, V, Liuyang Wang, Ken Luzinski, Petra Dufkova, Martina M. Vyskočilová, Michael W. Nachman, Pavel Munclinger, Miloš Macholán, Jaroslav Piálek, and Priscilla K. Tucker. 2012. “GenomeWide Architecture of Reproductive Isolation in a Naturally Occurring Hybrid Zone between Mus Musculus Musculus and M. M. Domesticus.” Molecular Ecology 21 (12): 3032–47. doi:10.1111/j.1365-294X.2012.05583.x.

Jones, ELEANOR P., JEROEN Van Der KOOIJ, ROAR Solheim, and JEREMY B. Searle. 2010. “Norwegian House Mice (Mus Musculus Musculus/domesticus): Distributions, Routes of Colonization and Patterns of Hybridization.” Molecular Ecology 19 (23). Blackwell Publishing Ltd: 5252–64. doi:10.1111/j.1365-294X.2010.04874.x.

Kaessmann, Henrik, Nicolas Vinckenbosch, and Manyuan Long. 2009. “RNA-Based Gene Duplication: Mechanistic and Evolutionary Insights.” Nature Reviews Genetics 10 (1). Nature Publishing Group: 19–31. doi:10.1038/nrg2487.

Katju, Vaishali, and Ulfar Bergthorsson. 2013. “Copy-Number Changes in Evolution: Rates, Fitness Effects and Adaptive Significance.” Frontiers in Genetics 4 (January): 273. doi:10.3389/fgene.2013.00273.

Kazmi, Syed J, Stephanie J Byer, Jenell M Eckert, Amy N Turk, Richard P H Huijbregts, Nicole M Brossier, William E Grizzle, Fady M Mikhail, Kevin A Roth, and Steven L Carroll. 2013. “Transgenic Mice Overexpressing Neuregulin-1 Model Neurofibroma-Malignant Peripheral Nerve Sheath Tumor Progression and Implicate Specific Chromosomal Copy Number Variations in Tumorigenesis.” The American Journal of Pathology 182 (3): 646–67. doi:10.1016/j.ajpath.2012.11.017.

Keane, Thomas M, Leo Goodstadt, Petr Danecek, Michael A White, Kim Wong, Binnaz Yalcin, Andreas Heger, et al. 2011. “Mouse Genomic Variation and Its Effect on Phenotypes and Gene Regulation.” Nature 477 (7364): 289–94. doi:10.1038/nature10413.

Kondrashov, F. a. 2012. “Gene Duplication as a Mechanism of Genomic Adaptation to a Changing Environment.” Proceedings of the Royal Society B: Biological Sciences, no. September. doi:10.1098/rspb.2012.1108.

Kondrashov, Fyodor A, and Eugene V Koonin. 2004. “A Common Framework for Understanding the Origin of Genetic Dominance and Evolutionary Fates of Gene Duplications.” Trends in Genetics: TIG 20 (7): 287–90. doi:10.1016/j.tig.2004.05.001.

Kruuk, L. E. B., S. J. E. Baird, K. S. Gale, and N. H. Barton. 1999. “A Comparison of Multilocus Clines Maintained by Environmental Adaptation or by Selection Against Hybrids.” Genetics 153 (4): 1959–71. http://www.genetics.org/content/153/4/1959.short.

Lappalainen, Ilkka, John Lopez, Lisa Skipper, Timothy Hefferon, J Dylan Spalding, John Garner, Chao Chen, et al. 2013. “DbVar and DGVa: Public Archives for Genomic Structural Variation.” Nucleic Acids Research 41 (Database issue): D936–41. doi:10.1093/nar/gks1213.

Locke, M Elizabeth O, Maja Milojevic, Susan T Eitutis, Nisha Patel, Andrea E Wishart, Mark Daley, Kathleen A Hill, et al. 2015. “Genomic Copy Number Variation in Mus Musculus.” BMC Genomics 16 (1). BioMed Central: 497. doi:10.1186/s12864-015-1713-z.

Lynch, M. 2000. “The Evolutionary Fate and Consequences of Duplicate Genes.” Science 290 (5494): 1151–55. doi:10.1126/science.290.5494.1151.

Lynch, Michael, and Allan G. Force. 2000. “The Origin of Interspecific Genomic Incompatibility via Gene Duplication.” The American Naturalist 156 (6). The University of Chicago Press: 590–605. doi:10.1086/316992.

Macholán, M, Stuart J E Baird, Petra Dufková, Pavel Munclinger, Barbora Vošlajerová Bímová, and Jaroslav Piálek. 2011. “Assessing Multilocus Introgression Patterns: A Case Study on the Mouse X Chromosome in Central Europe.” Evolution 65: 1428–46. doi:10.1111/j.1558-5646.2011.01228.x.

Macholán, M, Stuart J E Baird, Pavel Munclinger, Petra Dufková, Barbora Bímová, and Jaroslav Piálek. 2008. “Genetic Conflict Outweighs Heterogametic Incompatibility in the Mouse Hybrid Zone?” BMC Evolutionary Biology 8 (1): 271. doi:10.1186/1471-2148-8-271.

Macholán, M, Pavel Munclinger, Monika Šugerková, Petra Dufková, Barbora Bímová, Eva Božíková, Jan Zima, and Jaroslav Piálek. 2007. “GENETIC ANALYSIS OF AUTOSOMAL AND X-LINKED MARKERS ACROSS A MOUSE HYBRID ZONE.” Evolution 61 (4). Blackwell Publishing Inc: 746–71. doi:10.1111/j.1558-5646.2007.00065.x.

Macholán, M., Stuart J E Baird, Pavel Munclinger, and J. Pialek. 2012. Evolution of the House Mouse. Cambridge University Press.

Macholán, M., Pavel Munclinger, Monika Sugerková, Petra Dufková, Barbora Bímová, Eva Bozíková, Jan Zima, and Jaroslav Piálek. 2007. “Genetic Analysis of Autosomal and X-Linked Markers across a Mouse Hybrid Zone.” Evolution; International Journal of Organic Evolution 61 (4): 746–71. doi:10.1111/j.1558-5646.2007.00065.x.

Mejía-Benítez, María A, Amélie Bonnefond, Loïc Yengo, Marlène Huyvaert, Aurélie Dechaume, Jesús Peralta-Romero, Miguel Klünder-Klünder, et al. 2015. “Beneficial Effect of a High Number of Copies of Salivary Amylase AMY 1 Gene on Obesity Risk in Mexican Children.” Diabetologia 58 (2): 290–94. doi:10.1007/s00125-014-3441-3.

Nürnberger, Beate, Nick Barton, Catriona Maccallum, J. Gilchrist, and M. Appleby. 1995. “Natural Selection on Quantitative Traits in the Bombina Hybrid Zone.” Evolution 49 (6): 1224–38.

Otto, Sarah P., and Paul Yong. 2002. “The Evolution of Gene Duplicates.” In Advances in Genetics, 46:451–83. doi:10.1016/S0065-2660(02)46017-8.

Ottolini, Barbara, Michael J Hornsby, Razan Abujaber, Jacqueline A L MacArthur, Richard M Badge, Trude Schwarzacher, Donna G Albertson, Charles L Bevins, Jay V Solnick, and Edward J Hollox. 2014. “Evidence of Convergent Evolution in Humans and Macaques Supports an Adaptive Role for Copy Number Variation of the β-Defensin-2 Gene.” Genome Biology and Evolution 6 (11): 3025–38. doi:10.1093/gbe/evu236.

Paudel, Yogesh, Ole Madsen, Hendrik-Jan Megens, Laurent A F Frantz, Mirte Bosse, Richard P M A Crooijmans, and Martien A M Groenen. 2015. “Copy Number Variation in the Speciation of Pigs: A Possible Prominent Role for Olfactory Receptors.” BMC Genomics 16 (1): 330. doi:10.1186/s12864-015-1449-9.

Perry, George H, Nathaniel J Dominy, Katrina G Claw, Arthur S Lee, Heike Fiegler, Richard Redon, John Werner, et al. 2007. “Diet and the Evolution of Human Amylase Gene Copy Number Variation.” Nature Genetics 39 (10): 1256–60. doi:10.1038/ng2123.

Pezer, Željka, Bettina Harr, Meike Teschke, Hiba Babiker, and Diethard Tautz. 2015. “Divergence Patterns of Genic Copy Number Variation in Natural Populations of the House Mouse (Mus Musculus Domesticus) Reveal Three Conserved Genes with Major Population-Specific Expansions.” Genome Research 25 (8): 1114–24. doi:10.1101/gr.187187.114.

Pfennig, Karin S, and David W Pfennig. 2009. “Character Displacement: Ecological and Reproductive Responses to a Common Evolutionary Problem.” The Quarterly Review of Biology 84 (3). NIH Public Access: 253–76. http://www.ncbi.nlm.nih.gov/pubmed/19764283.

Phifer-Rixey, Megan, and Michael W Nachman. 2015. “Insights into Mammalian Biology from the Wild House Mouse *MusMusculus*” eLife 4 (April). doi:10.7554/eLife.05959.

Pirooznia, Mehdi, Fernando S Goes, and Peter P Zandi. 2015. “Whole-Genome CNV Analysis: Advances in Computational Approaches.” Frontiers in Genetics 6 (January): 138. doi:10.3389/fgene.2015.00138.

Quinlan, A. R., R. A. Clark, S. Sokolova, M. L. Leibowitz, Y. Zhang, M. E. Hurles, J. C. Mell, and I. M. Hall. 2010. “Genome-Wide Mapping and Assembly of Structural Variant Breakpoints in the Mouse Genome.” Genome Research 20 (5). Cold Spring Harbor Laboratory Press: 623–35. doi:10.1101/gr.102970.109.

Rajabi-Maham, H., A. Orth, and F. Bonhomme. 2008. “Phylogeography and Postglacial Expansion of Mus Musculus Domesticus Inferred from Mitochondrial DNA Coalescent, from Iran to Europe.” Molecular Ecology 17 (2): 627–41. doi:10.1111/j.1365-294X.2007.03601.x.

Redon, Richard, Shumpei Ishikawa, Karen R Fitch, Lars Feuk, George H Perry, T Daniel Andrews, Heike Fiegler, et al. 2006. “Global Variation in Copy Number in the Human Genome.” Nature 444 (7118): 444–54. doi:10.1038/nature05329.

Roberts, Chrissy H, Wei Jiang, Jyothi Jayaraman, John Trowsdale, Martin J Holland, and James A Traherne. 2014. “Killer-Cell Immunoglobulin-like Receptor Gene Linkage and Copy Number Variation Analysis by Droplet Digital PCR.” Genome Medicine 6 (3): 20. doi:10.1186/gm537.

Scavetta, Rick J., and Diethard Tautz. 2010. “Copy Number Changes of CNV Regions in Intersubspecific Crosses of the House Mouse.” Molecular Biology and Evolution 27 (8): 1845–56. doi:10.1093/molbev/msq064.

Schrider, Daniel R, and Matthew W Hahn. 2010. “Lower Linkage Disequilibrium at CNVs Is due to Both Recurrent Mutation and Transposing Duplications.” Molecular Biology and Evolution 27 (1): 103–11. doi:10.1093/molbev/msp210.

Servedio, Maria R, and Mohamed A F Noor. 2003. “The Role Of Reinforcement in Speciation.” Annual Review of Ecology, Evolution, and Systematics 34: 339–64.

Smadja, C., and G. Ganem. 2005. “Asymmetrical Reproductive Character Displacement in the House Mouse.” Journal of Evolutionary Biology 18 (6). Blackwell Science Ltd: 1485–93. doi:10.1111/j.1420-9101.2005.00944.x.

Smadja, C. M., Etienne Loire, Pierre Caminade, Marios Thoma, Yasmin Latour, Camille Roux, Michaela Thoss, Dustin J. Penn, Guila Ganem, and Pierre Boursot. 2015. “Seeking Signatures of Reinforcement at the Genetic Level: A Hitchhiking Mapping and Candidate Gene Approach in the House Mouse.” Molecular Ecology 24 (16): 4222–37. doi:10.1111/mec.13301.

Sokal, RR, and FJ Rohlf. 1995. Biometry. 3rd ed. New York: W.H. Freeman and Company.

Talley, H. M., Christina M. Laukaitis, and Robert C. Karn. 2007. “FEMALE PREFERENCE FOR MALE SALIVA: IMPLICATIONS FOR SEXUAL ISOLATION OF MUS MUSCULUS SUBSPECIES.” Evolution 55 (3): 631–34. doi:10.1111/j.0014-3820.2001.tb00796.x.

Turner, L M, D J Schwahn, and B. Harr. 2012. “Reduced Male Fertility Is Common but Highly Variable in Form and Severity in a Natural House Mouse Hybrid Zone.” Evolution; International Journal of Organic Evolution 66 (2): 443–58. doi:10.1111/j.1558-5646.2011.01445.x.

Wong, Kim, Suzannah Bumpstead, Louise Van Der Weyden, Laura G Reinholdt, Laurens G Wilming, David J Adams, and Thomas M Keane. 2012. “Sequencing and Characterization of the FVB/NJ Mouse Genome.” Genome Biology 13 (8): R72. doi:10.1186/gb-2012-13-8-r72.

Yalcin, Binnaz, Kim Wong, Amarjit Bhomra, Martin Goodson, Thomas M Keane, David J Adams, and Jonathan Flint. 2012. “The Fine-Scale Architecture of Structural Variants in 17 Mouse Genomes.” Genome Biology 13 (3): R18. doi:10.1186/gb-2012-13-3-r18.

Yanchukov, Alexey, and Stephen Proulx. 2012. “Invasion of Gene Duplication through Masking for Maladaptive Gene Flow.” Evolution 66 (5): 1543–55.

Yanchukov, Alexey, and Stephen R. Proulx. 2014. “Migration-Selection Balance at Multiple Loci and Selection on Dominance and Recombination.” PLoS ONE 9 (2): e88651. doi:10.1371/journal.pone.0088651.

Yeaman, Sam, and Sarah P Otto. 2011. “ESTABLISHMENT AND MAINTENANCE OF ADAPTIVE GENETIC DIVERGENCE UNDER MIGRATION, SELECTION, AND DRIFT.” Evolution 65 (7): 2123–29. doi:10.1111/j.1558-5646.2011.01277.x.

Yeaman, Sam, and Michael C. Whitlock. 2011. “THE GENETIC ARCHITECTURE OF ADAPTATION UNDER MIGRATION-SELECTION BALANCE.” Evolution 65 (7). Blackwell Publishing Inc: 1897–1911. doi:10.1111/j.1558-5646.2011.01269.x.

Yang, H., Y. Ding, L. N. Hutchins, J. Szatkiewicz, T. A. Bell, B. J. Paigen, J. H. Graber, F. Pardo-Manuel de Villena, and G. A. Churchill. 2009. A customized and versatile high-density genotyping array for the mouse. Nature Methods 6 (9):663–666.

