## Supplementary Information for "Copy number variation of the putative speciation genes in the European house mouse hybrid zone"

Supplemental Table 1. Assays used for ddPCR

| Assay Name | Amplicon Location(s) (mm10 assembly) | Primer 1 (5' -> 3') | Primer 2 (5' -> 3') | Probe (5' -> 3') |
|---|---|---|---|---|
| *** | singleplex & multiplex | | | |
| *Cma1* | chr14:55943671-55943757 | ATT AAG GAT AAG CAG CGC CTT G | TGC AGT GGC TTC CTG ATA AGA | /56-FAM/CAG TGA GCT /ZEN/GCA GTC AGC ACA A/3IABkFQ/ |
| *Defb7* | chr8:19497563-19497641 | AAC GAG CTT GCT ATC GGG AA | ACT TGC AGC ATT TGA AAC GAA | /56-FAM/GCG AAT GCC /ZEN/TGC AAC GGT GCA TTG GC/3IABkFQ/ |
| *Ifit3\|3b* | chr19:34587023-34587096<br>chr19:34611393-34611466 | TGC TCA TCA CAG TGA CCA TG | GAA ATG GCA CTT CAG CTG TG | /56-FAM/CCG CCA CAG /ZEN/TGA GGT CAA CCG GGA/3IABkFQ/ |
| *Smok2a* | chr17:13226029-13226090 | TGA TTT TGG ACT TGG CAT CC | CCT CAG GAG CAC TAA ATG GGT A | /56-FAM/CCA GGG CAA /ZEN/AAA CTA AAC TTA TTC TGT GG/3IABkFQ/ |
| *Speer4a* | chr5:26034084-26034196 | GAC CCA TGA CCT TTC CCC AG | AAA TCA GGA AGG CCA GAG CC | /56-FAM/ACG GCT CTG /ZEN/ACC CTT TCA GGC CAG AAC C/3IABkFQ/ |
| *Tert* | chr13:73627688-73627805 | CCT CTG TGT CCG CTA GTT ACA | TCT TTG TAC CTC GAG ATG GCA | /5HEX/CCC GTG GGC /ZEN/AGG AAT TTC ACT A/3IABkFQ/ |
| *Vmn2r5:6* | chr3:64416315-64416454 | ACT TGA TGT GCA GTT GTG GA | ACA TCA TGC TCA GCT TCT CC | /56-FAM/AGG GCA AAG /ZEN/GGA CTG GGA GGG ACC T/3IABkFQ/ |
| *Vmn2r6:7* | chr3:64584994-64585064 | CAG CTG GGT TGT GTG AAA TG | CTG CAA AAG GAG AAC CAG GA | /56-FAM/TGC CAG CGA /ZEN/AAA CGC ACA CAG AAC ACC/3IABkFQ/ |
| *** | only singleplex | | | |

| | | | | |
|---|---|---|---|---|
| *Plac9_1* | chr14: 25888287-25888413<br>chr14: 26028057-26028183<br>chr14: 26167671-26167797 | GAA AGT TGG AAG TGC GGT TG | ATT TGG ACC CAT CTG CTA GC | n/a |
| *Plac9_2* | chr14: 25891235-25891380<br>chr14: 26031003-26031148<br>chr14: 26170617-26170762 | TGG AGA ATG TGG AGG AAT GC | TGG TAA GCA GCA ACT CAG AC | |
| *Plac9_3* | chr14: 25894523-25894626<br>chr14: 26034292-26034395<br>chr14: 26173906-26174009 | AGC ACA GCA TAG GAA GGT TG | TTG CTC AAG ATA CCC ACA GC | |
| *Tex35* | Chr1: 157103435-157103575 | TCC TGT GTG AAG AAG CCA AG | TGT GTG GAT GTA GGT GCA TG | n/a |
| *Spaca4* | Chr7: 45725382-45725519 | GCA GCC TTT GTT GAT GAT GG | ATC AAG GAC TGC GTC TTC TG | n/a |
| *Defb8* | Chr8: 19445866-19445954 | GCA GCA TTT GAA AGG AGA TCC AC | CTT ACA TTC GAA ACG GAG GCA | n/a |

Supplemental Figure 1. Quantasoft © 1D signal profiles of the ddPCR assays

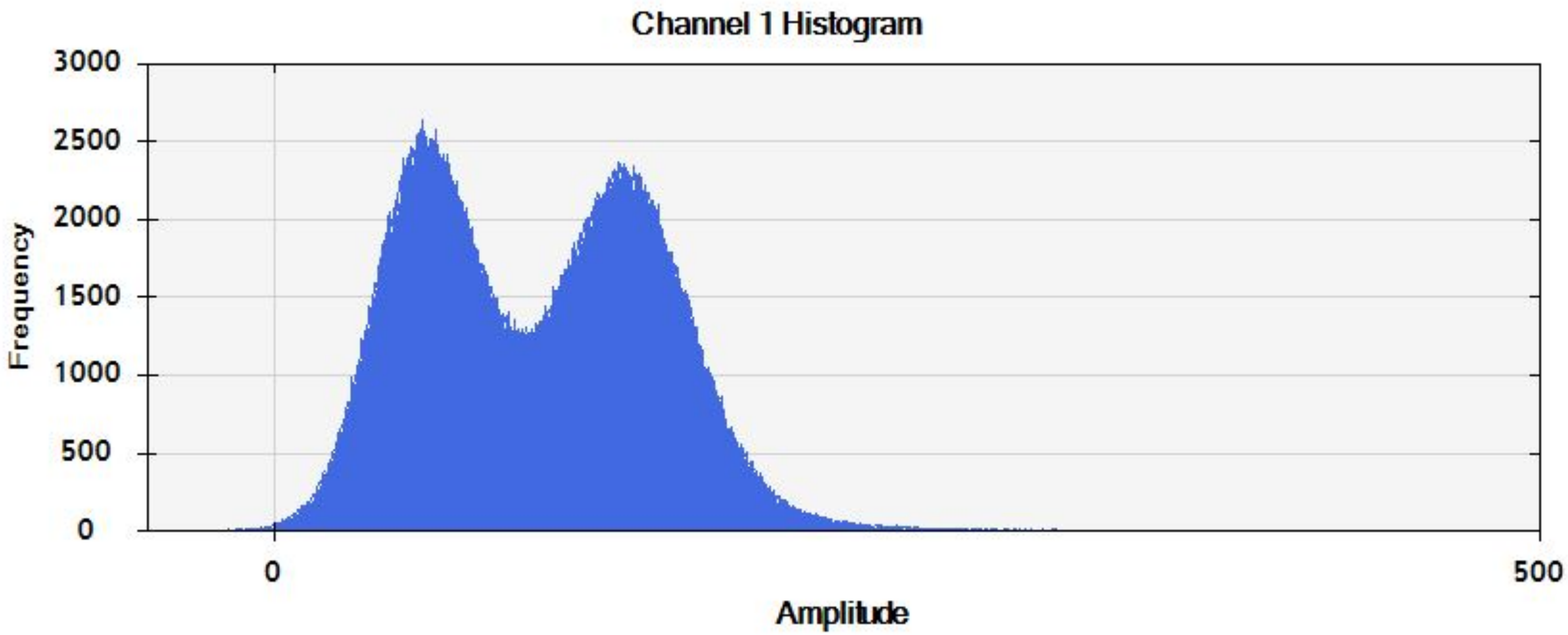
Channel 1 Histogram
3000
2500
2000
1500
1000
500
0
Frequency
0
500
Amplitude

*Defb7*

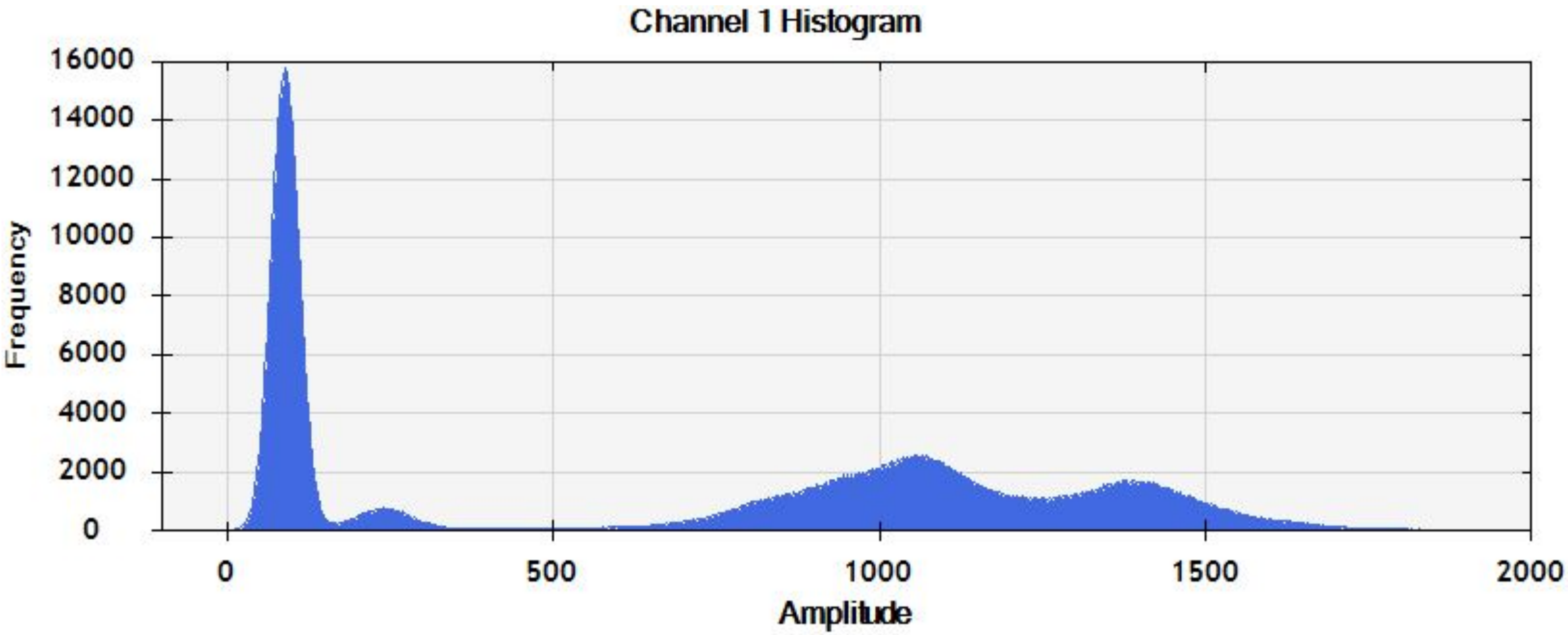
Channel 1 Histogram
Frequency
16000
14000
12000
10000
8000
6000
4000
2000
0
0
500
1000
1500
2000
Amplitude

*Ifit3/3d*

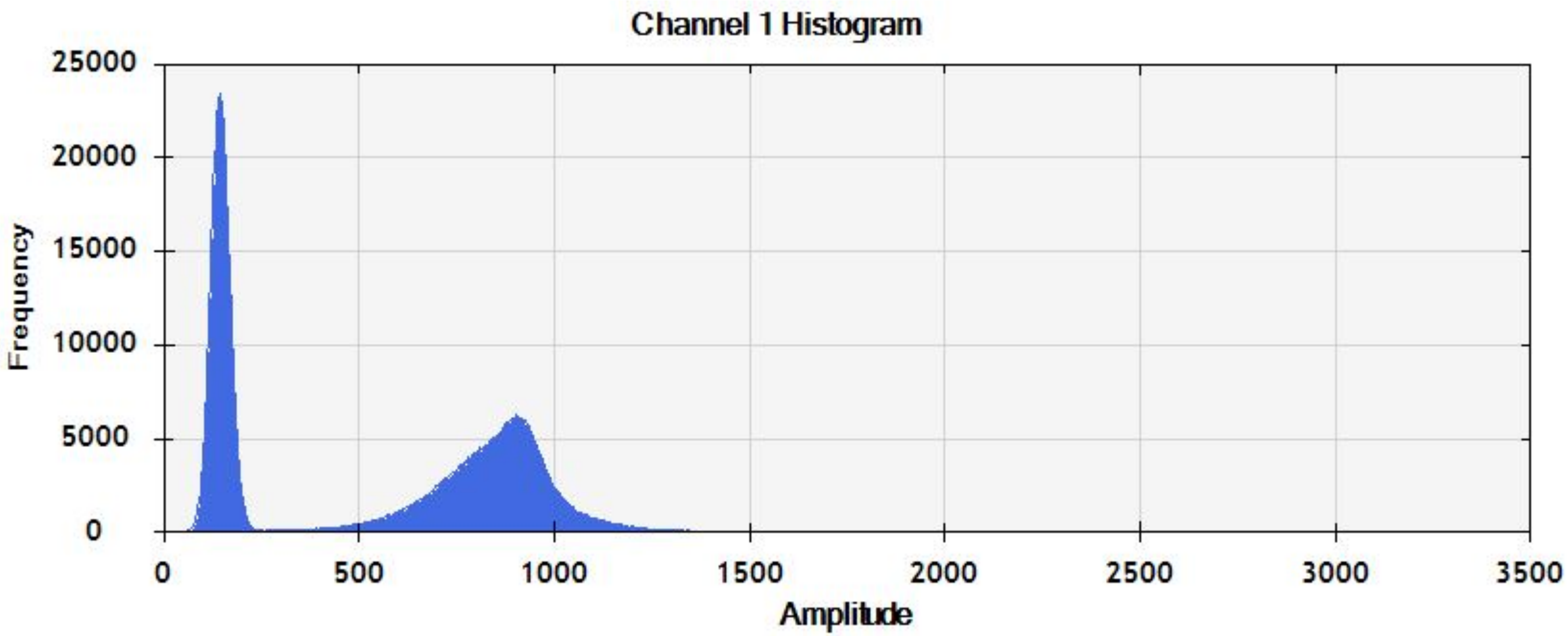
Channel 1 Histogram
25000
20000
15000
10000
5000
0
Frequency
0
500
1000
1500
2000
2500
3000
3500
Amplitude

***Vmn2r5:6***

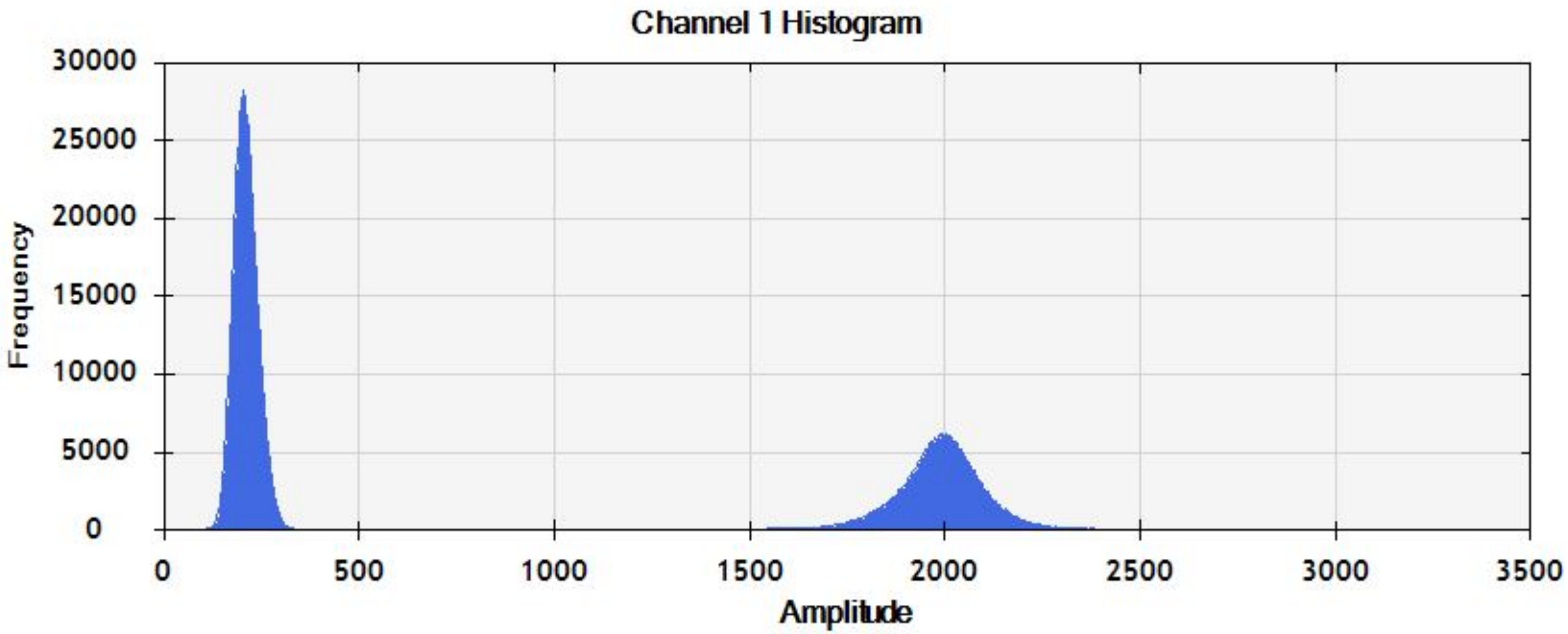
Channel 1 Histogram
Frequency
30000
25000
20000
15000
10000
5000
0
0
500
1000
1500
2000
2500
3000
3500
Amplitude

*Vmn2r6:7*

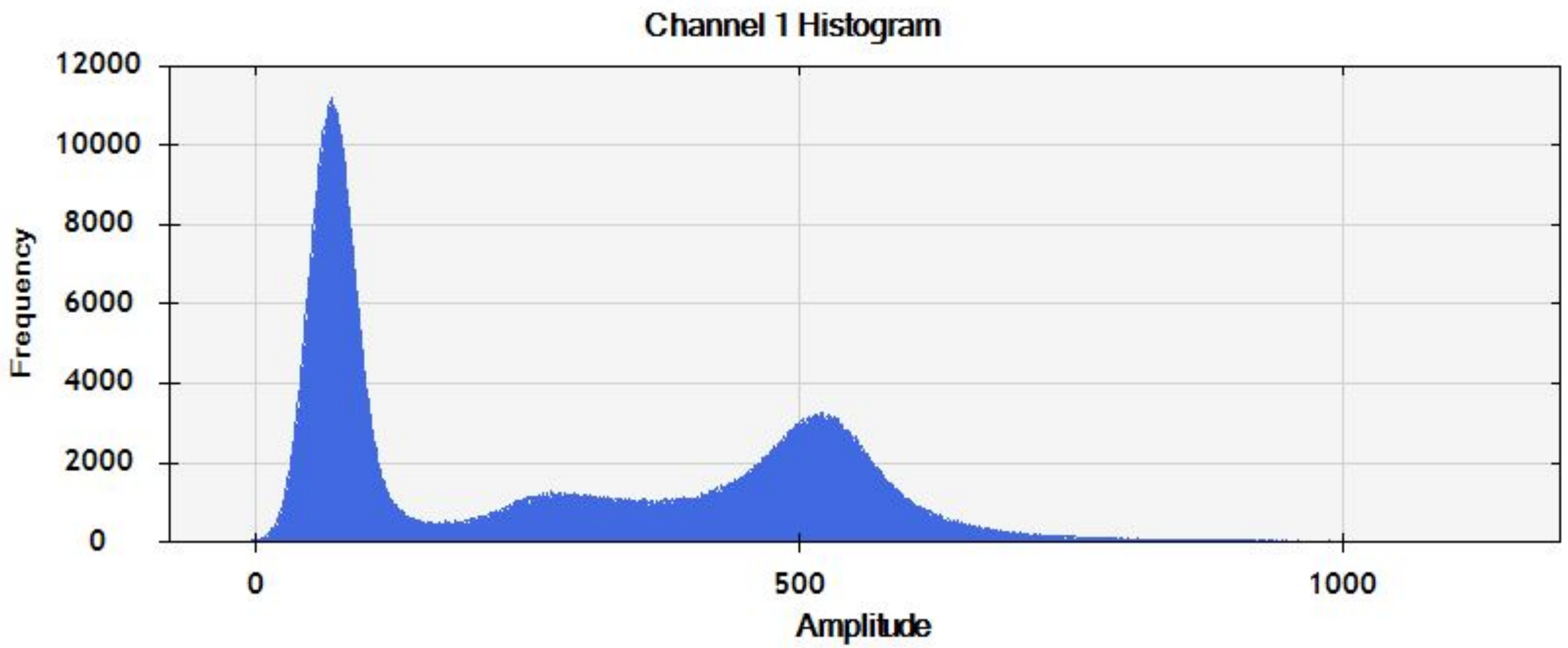
Channel 1 Histogram
12000
10000
8000
6000
4000
2000
0
Frequency
0
500
1000
Amplitude

***Speer4a***

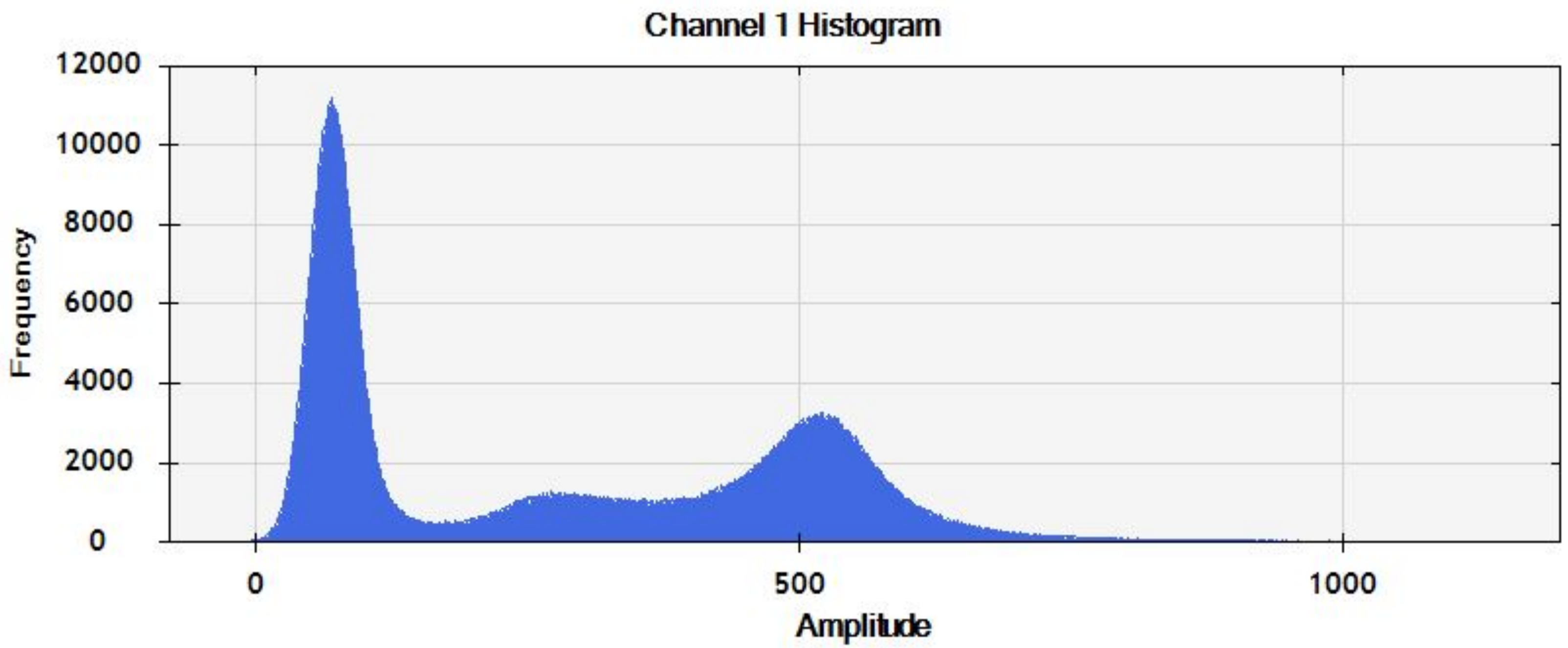

Channel 1 Histogram
12000
10000
8000
6000
4000
2000
0
Frequency
0
500
1000
Amplitude


***Smok2a***

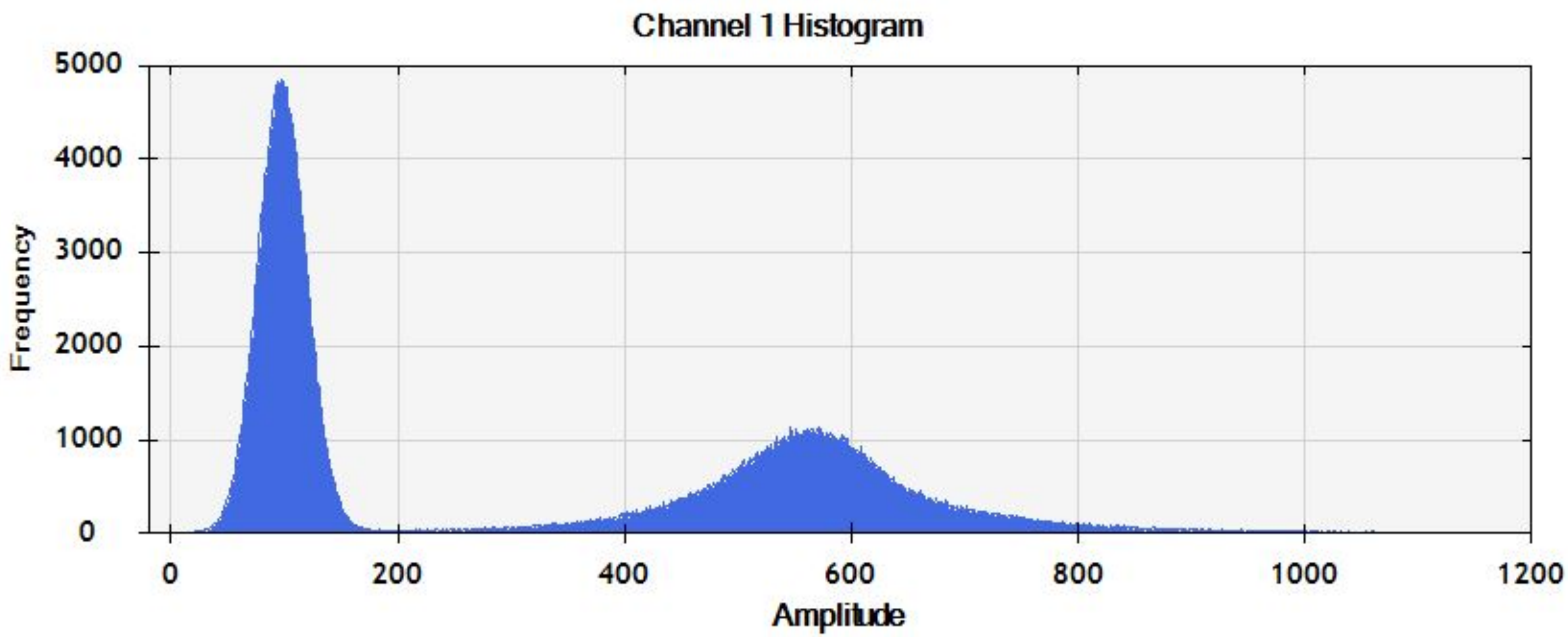

Channel 1 Histogram
5000
4000
3000
2000
1000
0
Frequency
0
200
400
600
800
1000
1200
Amplitude


***Cma1***

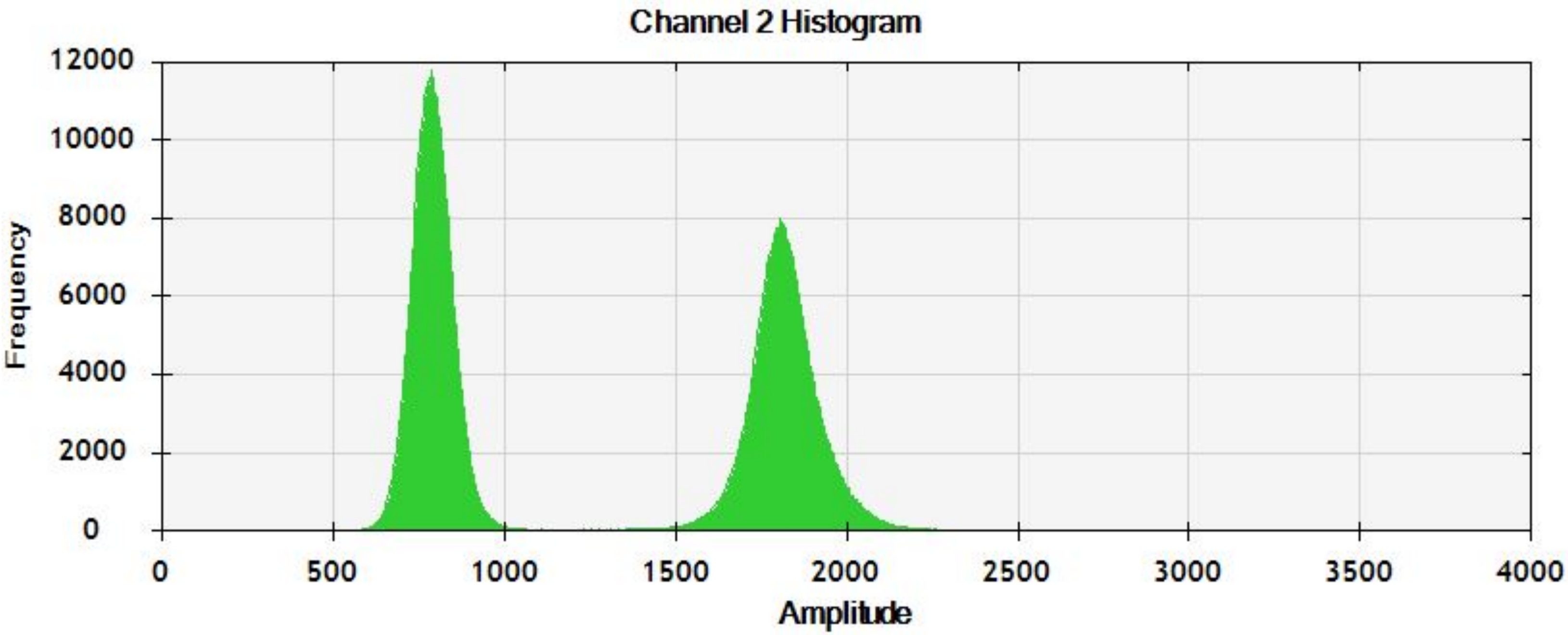
Channel 2 Histogram
Frequency
12000
10000
8000
6000
4000
2000
0
0
500
1000
1500
2000
2500
3000
3500
4000
Amplitude

***Tert***

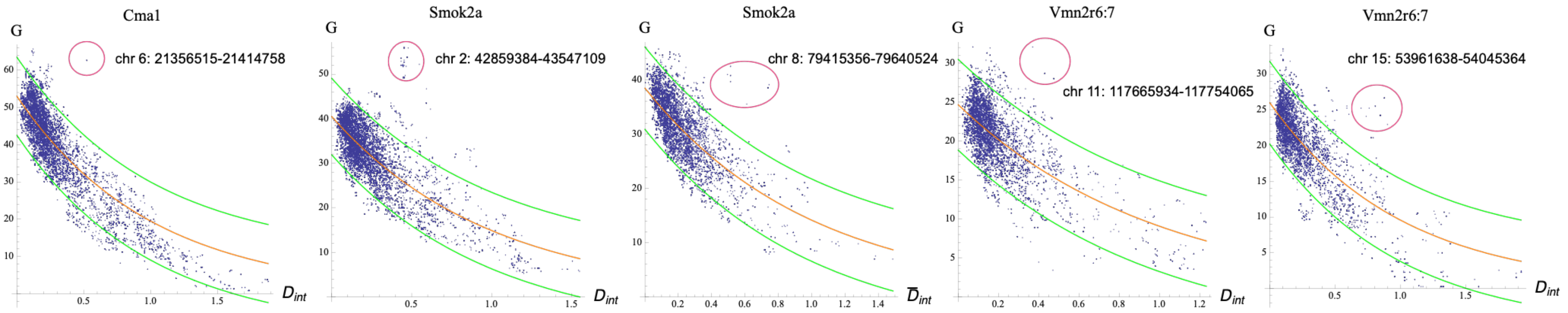


**Supplemental Figure 2.** Outliers in the chromosome-wide relationship between the per-SNP introgression index, $D_{\text{int}}$, and the [*mm-dd*] G-test (G).
